# Origins and breadth of pairwise epistasis in an α-helix of β-lactamase TEM-1

**DOI:** 10.1101/2021.11.29.470435

**Authors:** André Birgy, Clément Roussel, Harry Kemble, Jimmy Mullaert, Karine Panigoni, Audrey Chapron, Jérémie Chatel, Mélanie Magnan, Hervé Jacquier, Simona Cocco, Rémi Monasson, Olivier Tenaillon

## Abstract

Epistasis affects genome evolution together with our ability to predict individual mutation effects. The mechanistic basis of epistasis remains, however, largely unknown. To quantify and better understand interactions between fitness-affecting mutations, we focus on a 11 amino-acid α-helix of the protein β-lactamase TEM-1, and build a comprehensive library of more than 15,000 double mutants. Analysis of the growth rates of these mutants shows pervasive epistasis, which can be largely explained by a non-linear two-state model, where inactivating, destabilizing, neutral, or stabilizing mutations additively contribute to the phenotype. Hence, most epistatic interactions can be predicted by a non-linear model informed by single-point mutational measurements only. Deviations from the two-state model are consistently found for few pairs of residues, in particular when they are in contact. This result, as well as single-point mutation parameters, can be quantitatively found back through direct-coupling-analysis-based statistical models inferred from homologous sequence data. Our results thus shed light on the existence and the origins of the multiple determinants of the epistatic landscape, even at the level of small structural components of a protein, and suggest that the corresponding constraints shape the entire β-lactamase family.

## Introduction

Sequences of the first proteins triggered the emergence of molecular evolution and bioinformatics in the 1960s (Hagen, 2000). Yet, more than 50 years later, despite a massive number of available protein sequences and a pressing demand from human genetic disease and synthetic biology, the prediction of nonsynonymous mutation effects remains a challenging task.

Nonetheless, over the last decade, two independent approaches have offered new perspectives on the study of nonsynonymous mutation effects. Experimentally, protein deep mutational scans, in which the impacts of all possible single amino acid changes in a protein are investigated, have gained momentum allowing to study not only single mutants but also multiple mutants (Fowler and Fields, 2014). At the bioinformatics level, massive protein databases have allowed using multiple sequence alignment to infer the amino acids that are tolerated or not at a site. Interestingly, experimental and data-driven approaches revealed immediately that mutation impact could vary with genetic background (Jacquier *et al*., 2013; Bank *et al*., 2015, 2016). It was for instance shown that as little as a single mutation could change quite drastically the impact of many other mutations throughout a protein (Bloom *et al*., 2005; Jacquier *et al*., 2013). These observations called for a more comprehensive understanding of mutations’ effects and especially of their interactions.

Epistasis refers to the context-dependency of mutation effects. In population genetics, pairwise epistasis refers more precisely to mutation interactions that translate in non-additivity of log-fitness effects. Epistasis between mutation A and B can be quantitatively estimated as the deviation between the observed log-fitness of the double mutants, AB, and the sum of the log-fitness of both individual mutations (A and B) (Figure 1 (a)). Under this strict definition, epistasis has been predicted to impact significantly many facets of evolution, from the evolution of mutation rate and recombination (de Visser and Elena, 2007), to the diversity of adaptive path and the repeatability of adaptation (de Visser and Krug, 2014). These undoubtful significant consequences of epistasis now call for an integrated and mechanistic understanding of epistasis causes.

**Figure 1:**
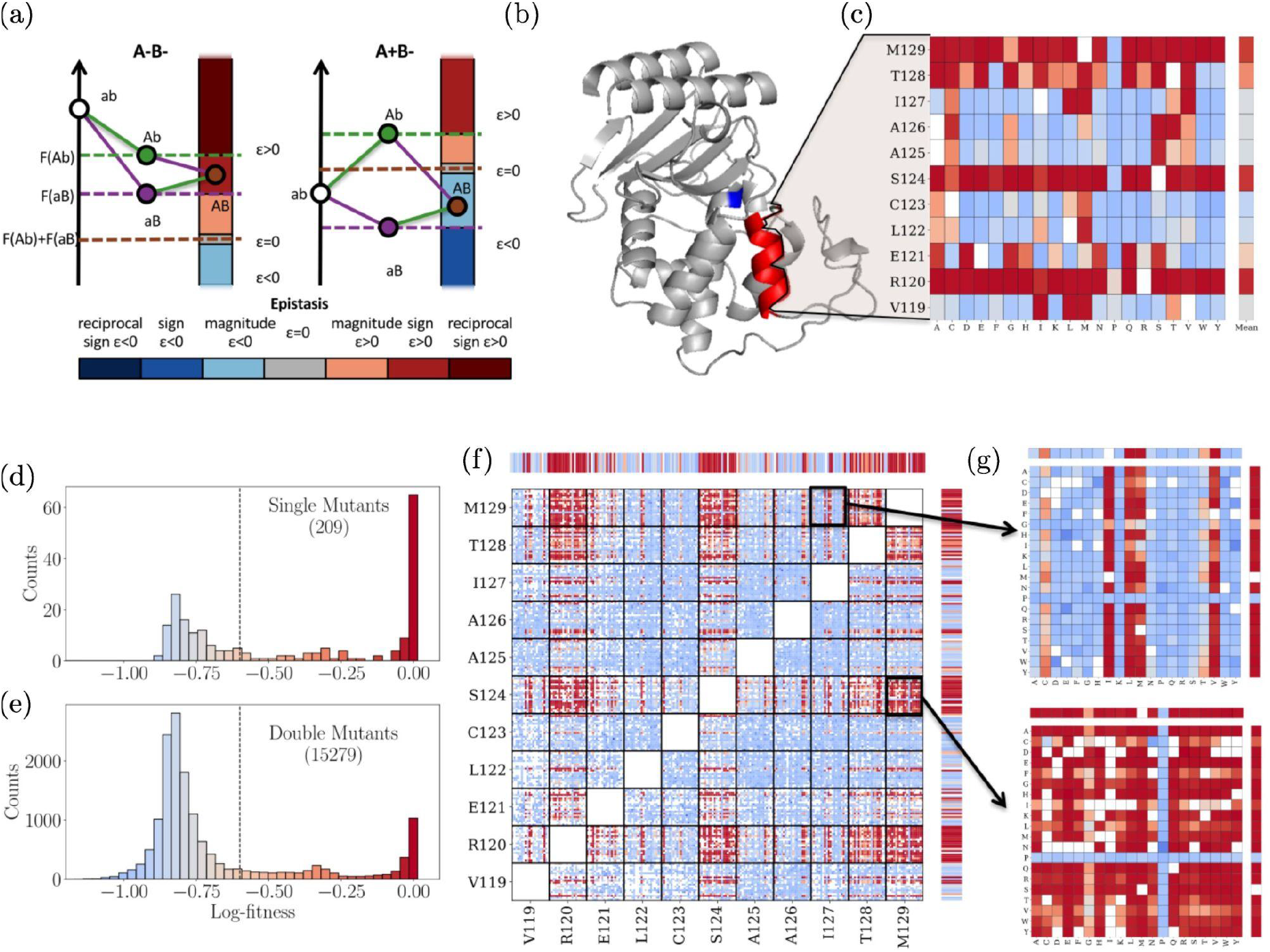
Single and double mutants’ log-fitness effects. (a) Pairwise epistasis measures the deviation of the observed log-fitness of a double mutant from the sum of the log-fitness of its single constituent mutants. It can also be qualitatively categorized as magnitude, sign, and reciprocal sign as well as positive or negative. The figures illustrate how this categorization functions in the case of a pair of deleterious mutations on the left and a pair including a deleterious (b to B) and a beneficial mutation (a to A) on the right (b) 3D structure of β-lactamase TEM-1. In red the α-helix of interest, and in blue the Serine residue of the active site. (c) The effects on the log-fitness of all single mutants per residue. Color scale is given in panels (d-e). (d-e) Distribution of log-fitness effects. Below the dotted line, mutants are considered non-functional. (d) For single mutants. (e) For double mutants. (f) Log-fitness of the double mutants with missing data in white. Color scale is given in panels (d-e). (g) Zoom on the double mutants log-fitness involving residues I127 and M129 on top and S124 and M129 at the bottom.

An integrated vision of epistasis may be obtained from a top-down perspective, with phenomenological models that capture its global properties. These models have shown that all forms of epistasis mentioned in Figure 1(a) can emerge from a simple nonlinear mapping of phenotype to fitness even if the phenotype is additive. For instance, all possible forms of pairwise epistasis are observed in the Fisher Geometric Model (Martin, Elena and Lenormand, 2007; Gros, Le Nagard and Tenaillon, 2009; Blanquart *et al*., 2014; Tenaillon, 2014), a smooth singled peaked phenotypic landscape in which fitness is a Gaussian function of the distance to an optimum phenotype. These observations motivated the research of an underlying simple phenotype that could explain globally the pattern of epistasis observed. Accordingly, statistical analysis of large datasets of multiple mutants have revealed epistasis to be largely described by an underlying additive phenotype (Otwinowski, McCandlish and Plotkin, 2018).

As proteins generally operate in a folded state, mutations’ impacts on protein have mainly been investigated through their effects on that fold or its affinity with a substrate. For epistatic interactions, two mutually non-exclusive mechanistic visions have emerged. With compensatory mutations, characterized by two independently deleterious mutations that, when combined, outcompete at least a single mutant, the idea of key-lock local interactions emerged. Alternatively, the existence of mutations with a global impact on protein stability (Bloom *et al*., 2005) hinted that the cooperative nature of protein stability could also result in epistatic effects, this time at a more global level (Wylie and Shakhnovich, 2011). The extent of both types of interactions and the overall prevalence of epistatic interactions remain however unclear.

To investigate the molecular determinant of epistatic interactions and our ability to predict it from a limited number of measures in a mutational scan, we generated a comprehensive library of more than 15,000 single and double mutants within an α-helix of β-lactamase TEM-1. TEM-1 is a highly successful antibiotic resistance gene present in about 35% of *Escherichia coli* natural isolates (EARS-Net France). We focused on an 11 amino acid α-helix, from residue 119 to 129 (Figure 1(b)), as α-helices are the most characterized and frequent secondary structure in protein folds. For the sake of generality, this α-helix is not involved in the active site; it is just a structural component of the enzyme. The mutants, which cover more than 76% of all possible double mutants, were analyzed for their impact on protein activity, measured through the minimum inhibitory concentration (MIC), and more importantly, through their effects on fitness, allowing a proper estimation of epistasis (Method). We then investigated how a simple biophysical two state model linking the sequence dependent phenotype to the fitness accounted or not for the observed epistasis. Finally, to validate the relevance of our measurement of epistasis and its mechanistic interpretation, we used the protein sequence of numerous distant homologues of TEM-1 to predict mutation effect and epistasis through Direct Couplings Analysis (DCA).

## Results

### Wide spread epistasis in a alpha helix

The distribution of log-fitness effects of single mutants had a bimodal structure with close to 50% lethal mutants (log-fitness < -0.6) (Figure 1(d)). This suggested an overall important role of that α-helix. The different residues had very different patterns, with four sites permissive to mutations, while the others were much more sensitive (Figure 1(c)). As expected, proline, which is known to be incompatible with α-helix structure (von Heijne, 1991), was lethal for the enzyme function or close to at all sites (log-fitness < -0.55) (Figure 1(c)). The distribution of double mutant effects appeared to be tri-modal with an even more significant fraction of loss of function genotypes (78%) (Figure 1(e)). A dominance effect emerged: mutant combinations including a lethal mutation were lethal. Out of the 10.887 double mutants involving at least a lethal mutant, only 105 (1.0%) had a log-fitness higher than -0.5 (Figure 2(a)). Only 2 (0.02% of total) resulted from the combination of two deleterious mutations, an instance of sign epistasis in which one of the mutations is deleterious in one background and beneficial in another. This general dominance effect clarifies the partial success of methods based on residue conservation (Ng and Henikoff, 2003; Adzhubei, Jordan and Sunyaev, 2013) to predict mutation effect: significant effects such as inserting a proline within an α-helix are effectively context-independent. This suggests that the key-lock epistatic compensations, characterized by two independently deleterious mutations that when combined outcompete at least a single mutant, are rare in the α-helix under study. We then focused on quantifying epistasis (Figure 2(a) and (b)) and noticed that double mutants’ log-fitness deviated substantially from the one expected, *i*.*e*. the sum of log-fitness of the two single mutants.

**Figure 2 :**
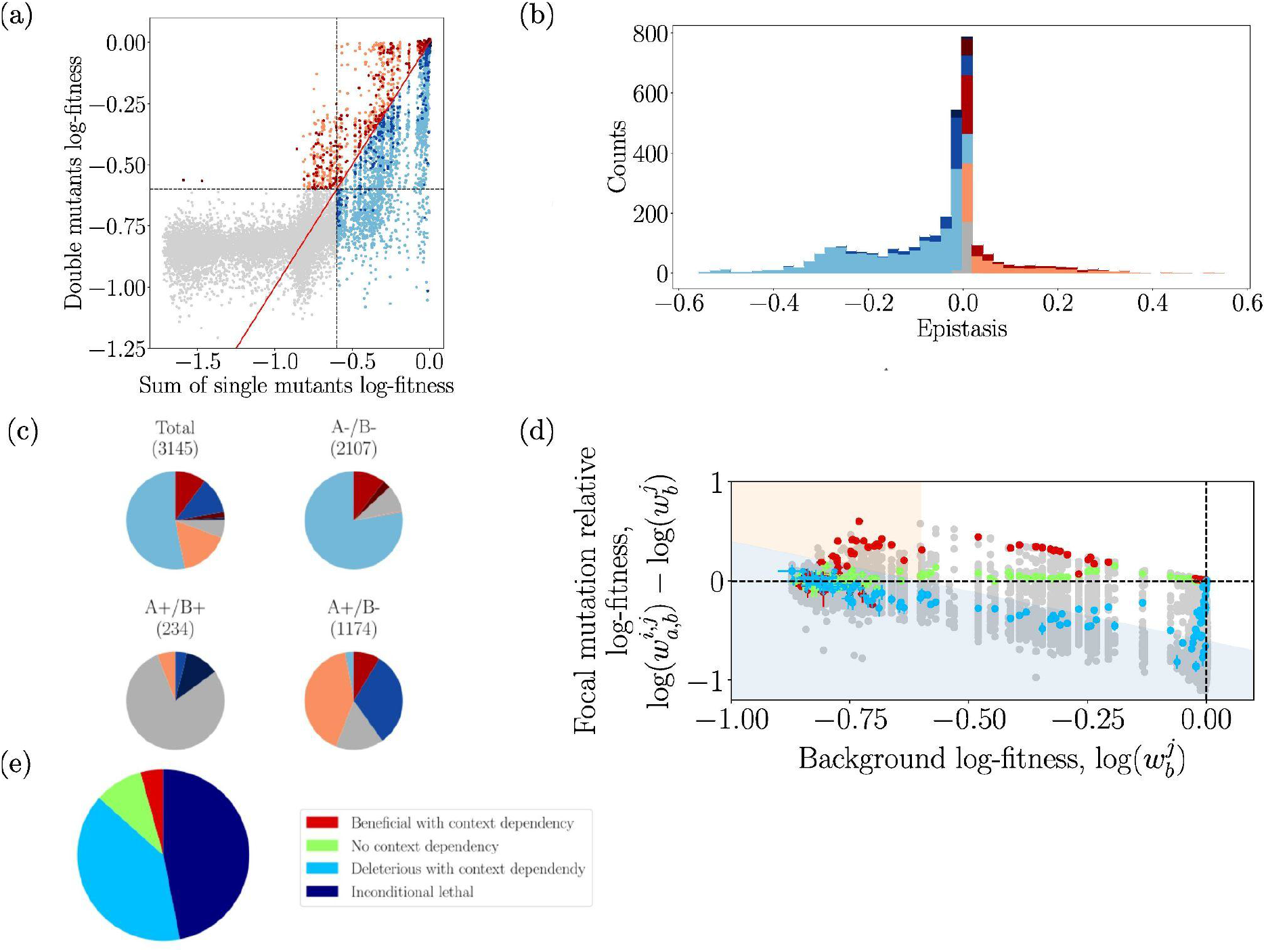
Pairwise epistasis. (a) Log-fitness of effects of double mutants, against the sum of the single mutants’ log-fitness. Grey mutants of observed log-fitness and predicted log-fitness based on single mutants lower than -0.6 cannot give reliable values for the epistasis. The colors of the other points represent the form of epistasis detected using the color code defined in Figure 1 (a). (b) Distribution of epistasis using the same color code, excluding mutants with non-measurable epistasis. (c) Categorization of epistasis for all mutations, pairs of deleterious (A-/B-), pairs involving one deleterious and one beneficial (A+/B-), or pairs of beneficial (A+/B+). (d) Relative log-fitness effect of all mutations against the log-fitness of the different backgrounds in which they were found. The values for three focal mutations, L122A, R120K, and S124E, are highlighted in blue, green, and red respectively. Blue shaded area corresponds to double mutants with fitness effects below the threshold, salmon shaded area corresponds to double mutant with log-fitness value higher than the threshold despite having a single mutant below it. (e) The fraction of mutations falling into unconditionally inactivating, deleterious with context-dependency, no context dependency, and beneficial with context-dependency is presented.

Epistasis could be estimated with high resolution only for non-lethal double mutants with non-lethal single mutants. Restricting the dataset to these mutants, we could compute a distribution of epistasis that was both broadly distributed around zero and biased towards negative values (Figure 2(b)), as observed on other experiments based on proxies of protein function rather than on true fitness, *i*.*e*. binding (Olson, Wu, Sun, 2014), or fluorescent (Sarkisyan *et al*., 2016). Yet, some instances of large positive epistasis were also found, especially among pairs including a beneficial mutation and a deleterious one (Figure 2(c)). We then looked at the log-fitness effect of individual mutations across all different backgrounds. For a given single mutant A, we plotted the log-fitness of the double mutants AB minus the log-fitness of the single mutant B (called focal mutation relative log-fitness) versus the log-fitness of single mutants B (called background log-fitness), see Figure 2(d). In this figure, the white area corresponds to mutants with high resolution on log-fitness for double mutants AB and single mutant B (log-fitness > -0.6). The blue region corresponds to lethal double mutants. And finally, the orange area corresponds to lethal single mutant B but where the double mutants AB have log-fitness greater than -0.6. Due to the high resolution of log-fitness in the white area, we are mainly interested in the patterns that exhibit the mutations in this area. These plots exhibit mutations with very contrasted and structured patterns that we grouped in four distinct categories (Figure 2 (e)).

Among the 209 possible single mutants, 98 (47%) are lethal across all backgrounds (the single mutants and al the double mutants including these single mutants having a log-fitness lower than -0.6). Due to the resolution of our experiments, we can not say so much about these loss of function mutations. Eighty three (40%) single mutants having log-fitness higher than -0.6 have negative epistasis, see for example blue points (Figure 2(d)). 19 (9%) mutations showed an overall context-independent mutation effect, *i*.*e*. have no epistasis, which correspond to a straight line with null slope in Figure 2(d), see for example green points (Figure 2 (d)). These mutants had a minor impact on log-fitness (less than a 1% effect on log-fitness). Finally, 9 (4%) mutations with marginally increased log-fitness effects in the ancestral background, *i*.*e*. have positive epistasis, see for example red points (Figure 2(d)).

Strikingly, excluding the 98 mutations that were lethal in all backgrounds, 83% of the mutations exhibited some strong form context dependencies that were structured by background log-fitness. A majority of double mutants AB associated with a given single mutant A exhibit either positive epistasis or null epistasis or negative epistasis, but not all three at once. This consistency suggests a macroscopic force at play, such as protein stability.

### A two-state model linking genotype to fitness is predictive of epistasis

One paradigm in protein analysis is that most residues in protein maintain the functional fold, and therefore mutations at these sites mainly alter its stability but not the activity (DePristo, Weinreich and Hartl, 2005). Protein stability has been described by a two-state model, corresponding to a functional folded state and to several nonfunctional unfolded states (Privalov, 1979; Wylie and Shakhnovich, 2011) (Figure 3(a) and (b)). The probability Pnat that a sequence correctly folds in its functional structure is

**Figure 3 :**
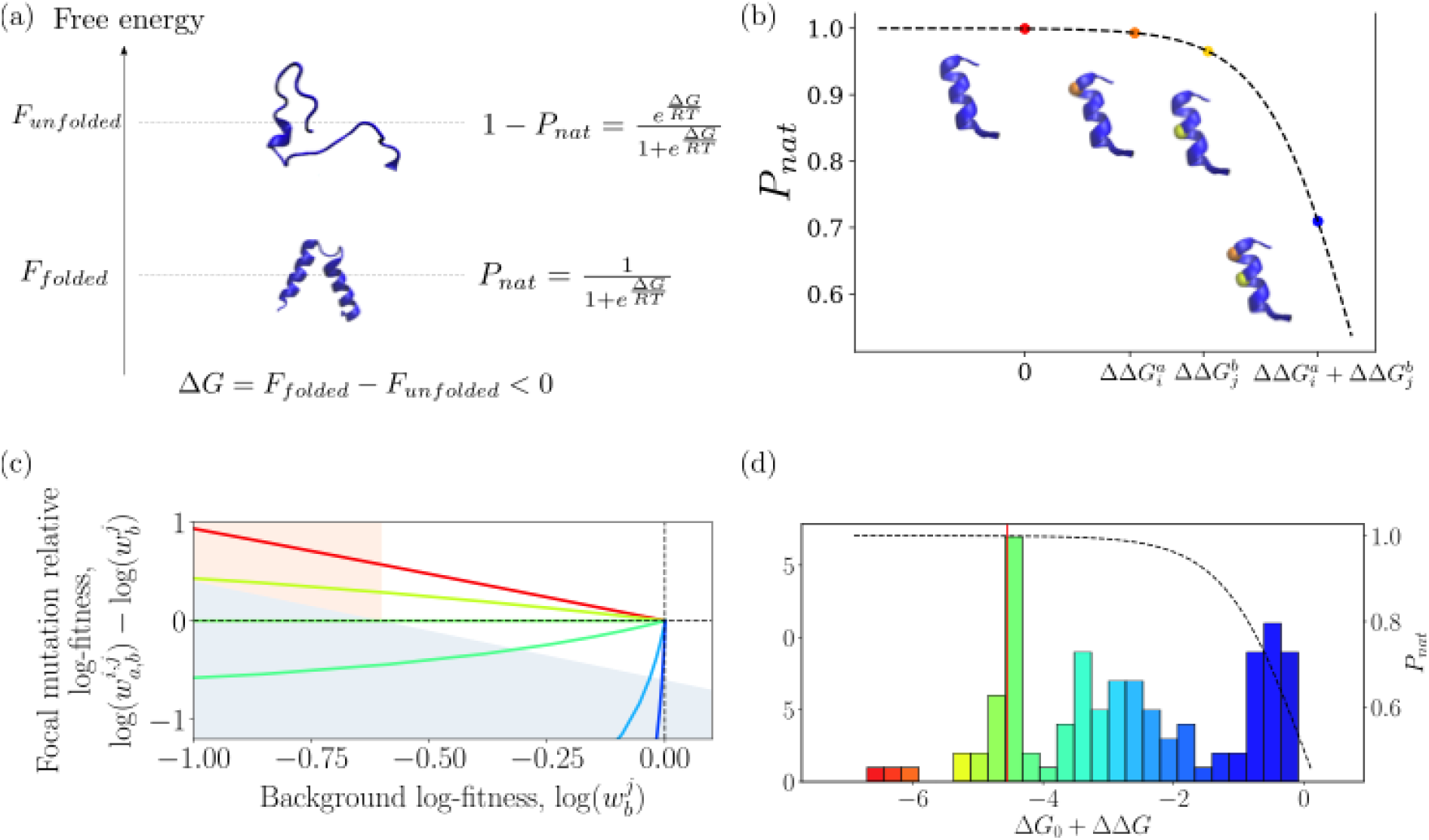
Stability and context-dependency. (a) Stability model. *P*_*nat*_ is the probability that the protein folds. (b) Effects of the mutations on the stability. Black dotted line corresponds to *P*_*nat*_. Red dot corresponds to the wild-type. Orange dot corresponds to a single mutation on the α-helix, with 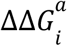. Yellow dot corresponds to a single mutation on the α-helix, with 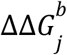 Blue dot corresponds to double mutations on the α-helix, with 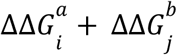. Mutations are considered as additive in ΔΔ*G*. However, this results in non-additive effect in *P*_*nat*_. (c) The relationship between background log-fitness and mutant’s relative log-fitness predicted by the model of stability is presented. The protein modeled has a free energy of -4.55 kcal mol^-1^, and the impact of mutations, ΔΔ*G*, is -2, -0.5, 0, 0.5, 2 and 3 from red to blue. (d) Histogram of the ΔΔ*G* estimated. Red line corresponds to Δ*G*_0_. Black dashed line corresponds to *P*_*nat*_ as a function of Δ*G*_0_ + ΔΔ*G*.

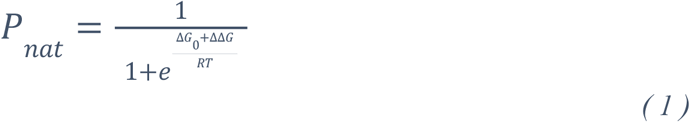

Upon change of the protein sequence, Pnat change according to the energetic impact of the mutations (ΔΔ*G*) on the free energy of the wild-type (Δ*G*_0_) stability.

One of the main hypotheses in this model is the additivity of the ΔΔ*G* upon multiple loci mutations:

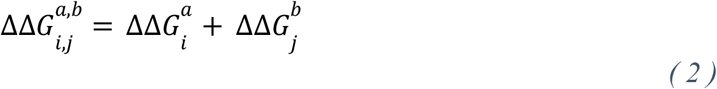

where 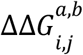 is associated with the double mutations at sites i and j with amino acids a and b, 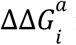 is associated with the single mutation at sites i with amino acids a, and 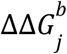 is associated with the single mutation at sites j with amino acids b.

We use the two-state model (1) here to more generally describe the sequence effect on the measured fitness, as previously done in (Wylie and Shakhnovich, 2011, Jacquier *et al* 2013, Otwinowski, McCandlish and Plotkin (2018)).

Therefore, the resulting log-fitness of a mutant can be computed as

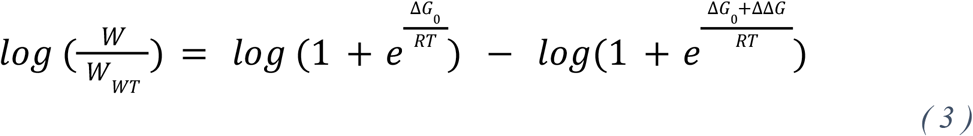

Depending on the mutant ΔΔ*G*, this model produces patterns of log-fitness effects according to background log-fitness similar to the one observed in the data (Figure 4 (a)). To have the best possible estimate of the parameters, we decided to estimate the ΔΔ*G* and Δ*G*_0_ from the log-fitness of single and double mutants (Method). As we accurately measure the log-fitness only above a threshold of -0.6, we keep only the 111 single mutants (53% of the total) with a log-fitness greater than -0.6. For each pair of previously chosen single mutants, the associated double mutant is kept if it has been measured experimentally: however its log-fitness is thresholded at -0.6. The stability model is itself thresholded at -0.6 during the inference. Keeping the thresholded lethal double mutants allows a better estimation of the ΔΔ*G*. We found Δ*G*_0_= -4.55 kcal.mol^−1^. To control experimental reproducibility, experiments have been carried out on two biological semi-replicates. For both biological semi-replicates, inferred ΔΔ*G* are highly correlated (r^2^ = 0.99, Supplementary Figure B(c)).

**Figure 4:**
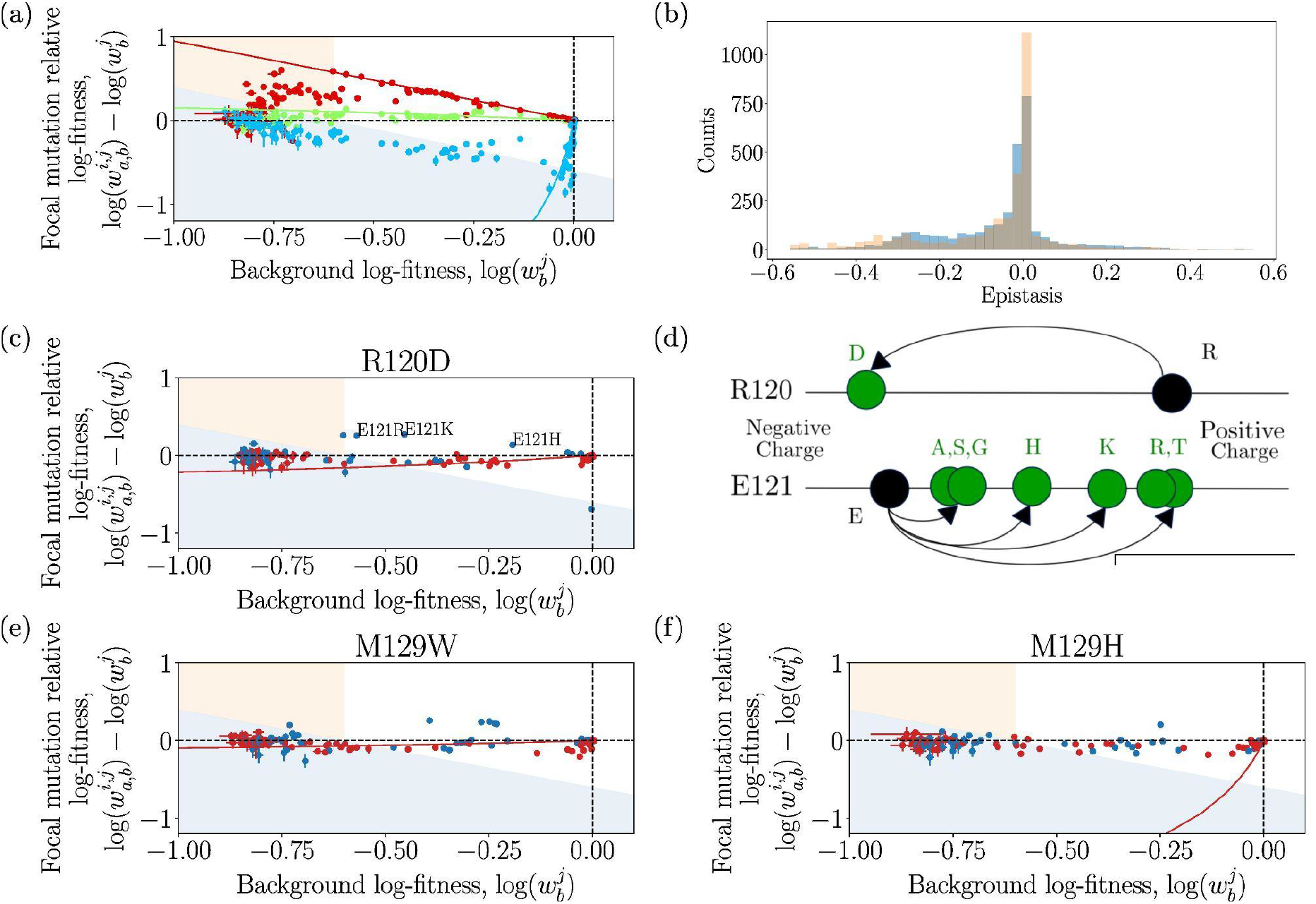
(a) The lines represent the fit of the model for the three mutants from Figure 3(d). Due to the resolution of our experiments, the lines are valid only in the white area. (b) In blue is the distribution of epistasis as presented in Figure 3(b), and overlaid on it in orange is the distribution of epistasis obtained with the fitted stability model. Deviations from the stability model. Relative log-fitness according to background fitness for three mutants: R120D (c), M129W (e), and M129H (f). Red dots represent distant sites and blue dots nearby sites. (d) At residue 120, the decrease of charge associated with R to D mutation compensates mutations at residue 121 that increased the charge.

As shown in Supplementary Figure C(a), the stability model well reproduces the experimental log-fitness of all the selected single (ρ = 0.99, r^2^ = 1.0) and double mutants (ρ = 0.92, r^2^ = 0.85). The above correlation is an improvement with respect to the one (ρ = 0.87, r^2^ = 0.65, Supplementary Figure C(b)) obtained by neglecting epistasis, *i*.*e*. assuming that the fitness of double mutations is the product of the ones of simple mutations. Moreover, as shown in Figure 4(a) the stability model captures the overall dependency of the fitness of a double mutant from the fitness of the background simple mutant. Finally, Figure 4 (b) shows that it reproduces the shape and breadth of the distribution of epistasis, with correlation ρ = 0.81, r^2^= 0.55 between observed and predicted epistasis. Our results are also consistent with previous experiments: R120G is known to have a stabilizing effect (Bershtein, Goldin and Tawfik, 2008; Salverda, De Visser and Barlow, 2010) and this effect is indeed captured by the model, with ΔΔG = -1.85 kcal.mol^−1^ (negative ΔΔ*G* corresponds to stabilizing mutation).

Supported by the ability of the two-state model to correctly capture the overall background dependency of the mutants and the epistasis, we then investigated its performance in predicting double mutation effects and epistasis when estimating its Δ*G*_0_ and ΔΔ*G* parameters on single mutations only. This amounts to reducing the number of mutational datapoints on which the model is built from 4493 to 111. As shown in Figure 5b, we find a remarkable Spearman correlation in epistatic predictions ranging from 0.6 to 0.7. Such dispersion of Spearman values comes from the lack of a precise determination on the estimation of Δ*G*_0_ arising from flat directions in the log-likelihood of the two-state model inferred only from single mutations (Supplementary). In particular as appears clearly when comparing Fig. 5b to Fig. 5a, the predictions related to positive epistasis are less accurate when estimating the parameters only from single mutations. The log-fitness of single mutants with stabilizing effect (ΔΔ*G* < 0) are close to 0 and inferring ΔΔ*G* only with single mutants lead to ΔΔ*G* = 0, while their stabilizing effect only appears when double mutants are taken into account.

**Figure 5:**
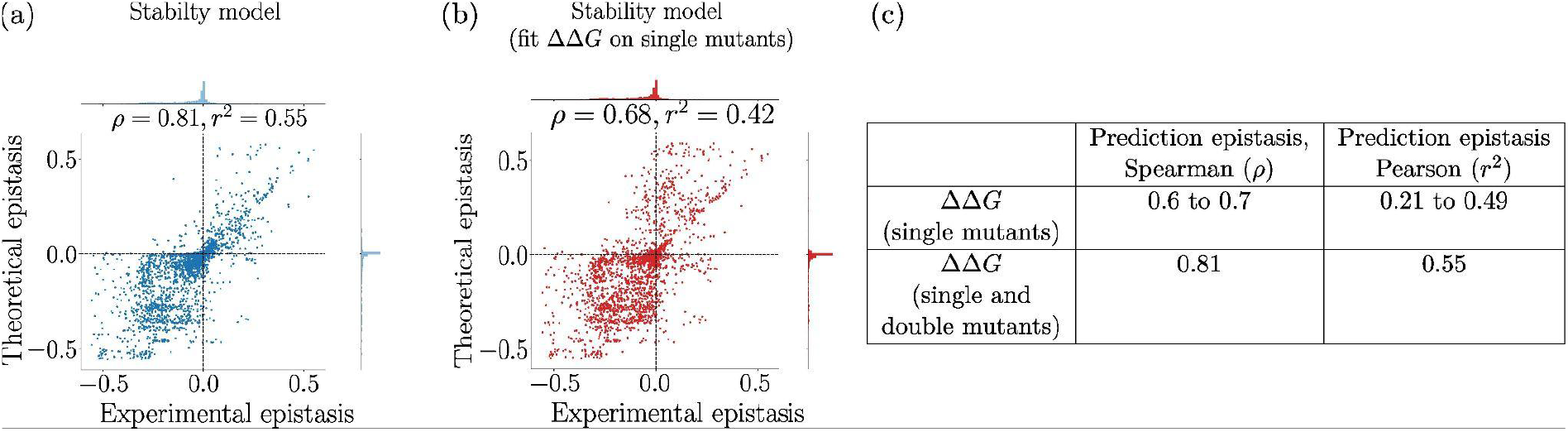
(a) Epistasis estimated from the two-state model fitted on single and double mutational data, against experimental epistasis. (b) Epistasis predicted with two-state model fitted only on single mutational data and fixed Δ*G*_0_ = -4.55 kcal.mol^-1^. (c) Correlations between predictions and experimental values for epistasis from two-state model with parameters ΔΔ*G* fitted only on single mutants by varying Δ*G*_0_ from -7 kcal.mol^−1^ to -3 kcal.mol^−1^, range corresponding to equivalent log-likelihood of the inferred model, compared to the full model where ΔΔ*G* and Δ*G*_0_ =-4.55 kcal.mol^−1^ are fixed on single and double mutants.

### Deviation from the two-state model are more frequent between physically close residues

While the two-state model is overall able to reproduce our data, we then investigated on which pairs of sites its predictions mostly deviated from them. First, when keeping only the residues at less than 6Å the correlation between experimental and predicted log-fitness with the two-state model decreased to ρ = 0.88, r^2^ = 0.80, while when only distant pairs (>6Å) were considered the correlation improved to ρ = 0.95, r^2^= 0.89 (Supplementary Figure C (c) and (d)). Accordingly, a maximum likelihood approach (Supplementary) estimated that the deviations to the two-state model were 1.28 times greater for close pairs of sites (<6Å) compared to distant ones (>6Å). This implies that our model explained less well the interactions between nearby sites than distant sites, suggesting that for local interactions, alternative forces could be at play.

To further characterize deviations from two-state model predictions, we computed, for each pair of residues i,j, the mean square error between the experimental log-fitness 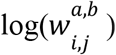 and the log-fitness predicted with the stability model 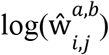 (Equation (3)),

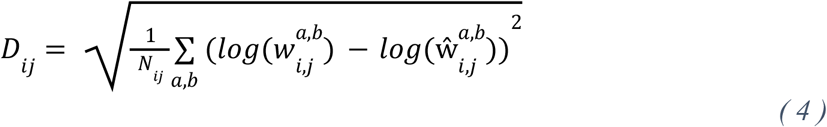

where *N*_*ij*_ is the total number of double mutants for which we can calculate the log-fitness according to the stability model for the pair i,j (Supplementary Figure D for the distribution of *D*_*ij*_). Large deviations *D*_*ij*_ imply that the assumption of linearity of the ΔΔ*G* embedded in the two-state model (Equation 2) is no longer valid for the corresponding pair. We refer to such pairwise interactions, not captured by the stability model, as idiosyncratic epistasis. For instance, mutation R120D and M129W showed signs of both positive and negative epistasis, the positive epistasis being restricted to residues in direct contact (Figure 4(c) and (e)). R120D mutation leads indeed to a change in charge, deleterious for distant interactions, which becomes beneficial when associated with departure from E121 charged amino acid, the neighboring amino acid (Figure 4(d)). The five pairs of sites with the largest idiosyncratic epistasis are: 128-129, 124-128, 123-127, 127-128 and 120-123. Among these five pairs, four correspond to the residues at less than 6Å, comforting that local interactions involve the largest deviation from the two-state model.

### Sequence of TEM-1 homologs can be used to predict mutation effects in TEM-1

At this stage the analysis of our experimental data suggests that epistasis results largely from a non-linear relationship between the sequence of a protein and its macroscopic fitness, well captured by a two-state model. Moreover, the deviations to the model are not random and occur preferentially for residues in contact, revealing this time some idiosyncratic epistasis. We next wanted to validate that these observations on epistasis, made by measuring fitness in the laboratory at a given antibiotic concentration, could be representative of generic properties of epistasis in the TEM-1 protein family, class A β-lactamases that evolved for millions of years.

To this aim we trained a model on Multiple Sequence Alignment (MSA) built on high-quality homologs of class A β-lactamases cleaned by hand (Philippon *et al*., 2016, 2019) and enriched on SwissProt and TrEMBL (The UniProt Consortium, 2021) (Method). The main idea is to learn a probability distribution over all the sequences **a** of length L from the MSA: sequences with high probability should correspond to putative β-lactamase. Each sequence **a** is supposed to be drawn from a Boltzmann distribution 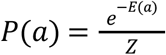. Once trained, we score all the single and double mutants according to their energies *E(****a***). For such model, known as Potts models, *E(****a***) reads

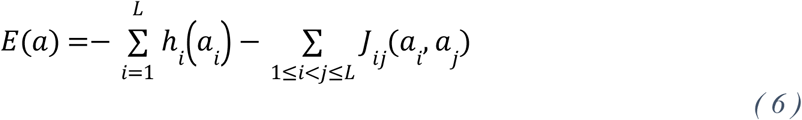

with fields *h*_*i*_(*a*_*i*_) and pairwise interactions *J*_*ij*_(*a*_*i*_, *a*_*j*_). Potts model, used in Direct-Couplings Analysis (DCA) (Weigt *et al*. 2009; Morcos *et al*. 2011) can disentangle direct coevolutionary couplings from indirect ones. DCA is widely used to predict tertiary contacts in proteins. This family of models was successfully used to design functional proteins with limited homology to existing sequences (Russ *et al*. 2020), and for TEM-1, they have been used for predicting fitness effects of single mutations (Figliuzzi *et al*. 2016).

To compare the predictions *E(****a***) and the results of the experiments, we need a proxy to link the two quantities. The most common proxy is the difference of log-likelihood between the mutant *a*_*mut*_ and the wild-type *a*_*WT*_ (Figliuzzi *et al*., 2016; Hopf *et al*., 2017; Zhao *et al*., 2021)

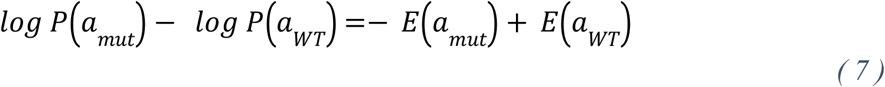

With that proxy Potts energies were found to be correlated to MIC (Figliuzzi *et al*., 2016), specificity constant *k*_*cat*_(Zhao *et al*., 2021), log-fitness (Hopf *et al*., 2017) or binding energies (Salinas and Ranganathan, 2018). Accordingly, we found a Spearman correlation ρ = 0.86 for the 209 single mutants and ρ = 0.64 for the 15.279 double mutants. This has to be compared with the comparable, but slightly worse results: ρ = 0.81 and ρ = 0.59, obtained with the independent model that only considers conservation of amino acids and not coevolution between sites (same Hamiltonian as Potts model but without couplings *J*_*ij*_ (*a*_*i*_, *a*_*j*_)). The Potts model, allows to better estimate the effects of the mutations, thanks to the couplings *J*_*ij*_ (*a*_*i*_, *a*_*j*_) which takes into account the background of TEM-1, instead of having an average global effect consistent across all the class A β-lactamases, as in the case of the independent model.

As shown in Figure 6(a) and (b)) the relation between the log-fitness and our Potts energy is highly nonlinear with a characteristic S-shape displaying saturation of the log-fitness both at large and small Potts energy values. As it is shown in Supplementary Figure F, the S-shape depends, as expected, on the experimental proxy to measure the fitness and changes when using the MIC instead of the relative enrichment at a fixed concentration, accordingly to the non-linear relation between MIC and relative enrichment (Supplementary Figure A(b)). For the independent model, the relationship is even more bimodal (Supplementary Figure E(a) and (b)). Due to the above nonlinear relation between the Potts model energy and the experimentally measured epistasis, the Potts model failed to predict epistasis (ρ = −0.06) when using directly the energy differences as epistasis proxy:

**Figure 6:**
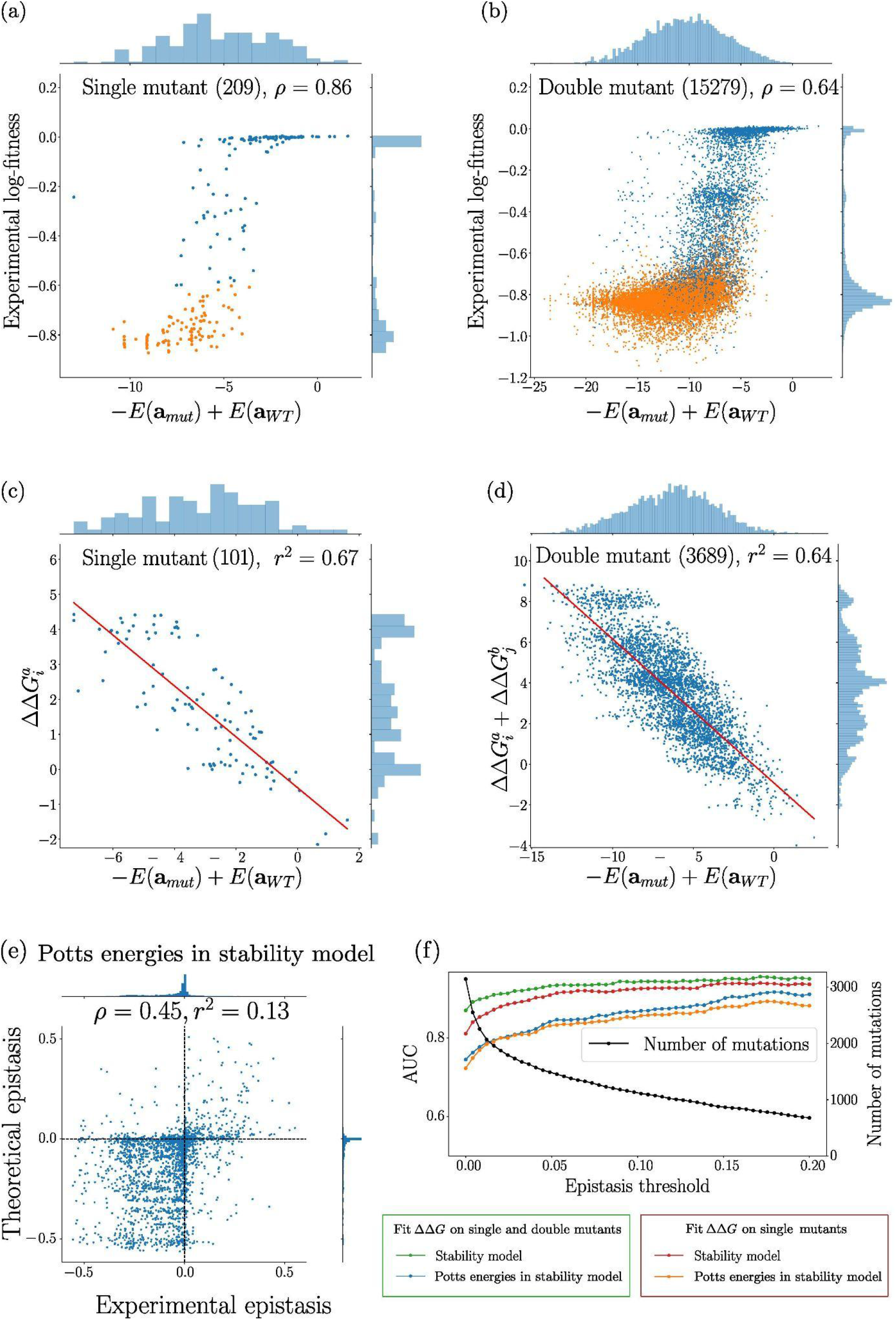
DCA Predictions. Energy from Potts model inferred from sequence data, versus experimental quantities. Blue dots correspond to mutations we used to estimateΔΔ*G*. Orange points are the other experimental mutations. For panels (c) and (d), only mutations present in the MSA are kept (with a different background than TEM-1), decreasing the number of single mutants from 111 to 101 and the number of double mutants from 4392 to 3689. (a)Experimental log-fitness against 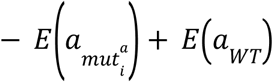 for single mutants. (b) Experimental log-fitness against 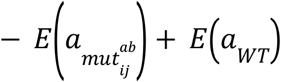 for double mutants. (c) 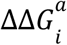 against 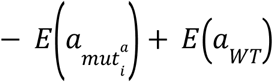 (d) 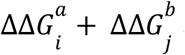 against 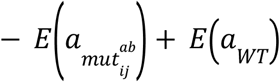. (e) Estimated epistasis with Potts energies in the stability model against experimental epistasis. Our predictions capture the sign of the experimental epistasis. (f) AUC against epistasis’ threshold for the different models. We used a threshold for the epistasis, keeping only experimental epistasis above this threshold in absolute value.

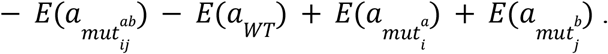

Yet, the typical “S” shape between the log-fitness and the Potts model energies is reminiscent of the relationship described by the two-state model. By directly comparing the Potts energies to the two-state model ΔΔ*G* parameters, related to the free energy changes due to the single mutations, in, we observed a much more linear relationship as shown in Figure 6(c) and 6(d) :

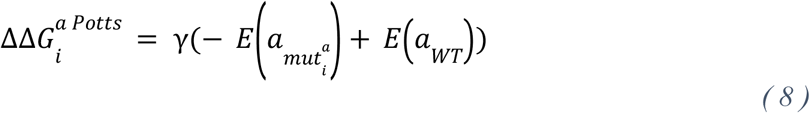

for single mutants and for double mutants we replace 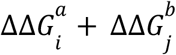 by:

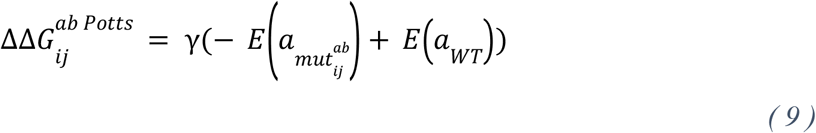

By restricting only to mutations that are present in the MSA and for which ΔΔG are estimated and by fitting as only parameter the slope γ = -0.71, we obtain a squared correlation coefficient r^2^ = 0.67 / 0.64 for single/double mutations respectively, between the Potts energies and the free energy parameters directly fitted from the mutational scan. The relation between ΔΔG and the energy of the independent model is less linear (r^2^ = 0.37 for the single mutants and r^2^ = 0.38 for the double mutants, Supplementary Figure E(c) and (d)), showing that the couplings *J*_*ij*_(*a*_*i*_, *a*_*j*_),which takes into account the TEM-1 background, are paramount to have an accurate estimation of the mutational effects from distant homologous sequences.

We have then used the two state model, Equation (3) with the previously fitted Δ*G*_0_ = -4.55 kcal.mol^−1^ and Potts free energy parameter 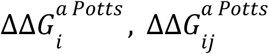 from *Eq*. (8, 9) to predict the log-fitness of single and double mutants, and the epistasis. In contrast to direct predictions above, now the predicted epistasis and the experimental one are correlated (ρ = 0.44), as shown in Figure 6(e). Figure 6(e) shows that the model principally captures the sign of the epistasis, as quantified by the AUC-ROC curve shown in Figure 6(f).

### Sequence of TEM-1 homologues predicts pairs of sites with idiosyncratic epistasis

As mentioned before, without encompassing the non-linearity of the two-state model, the Potts model fails to capture the epistatic effects (ρ = 0.06 between experimental values and predictions). Nevertheless, since Potts models are powerful tools to determine contacts between residues in protein structures, we investigated if these models could be predictive of the identified idiosyncratic epistatic interactions we detected. For Potts model, the canonical proxy to measure the interactions between two specific sites is the Frobenius norm of the couplings matrices 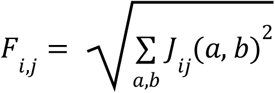, with the average product correction (Dunn, Wahl and Gloor, 2008). The top couplings of this metric are traditionally used to predict the tertiary contacts (Morcos *et al*., 2011).

We found that among the five pairs of sites with the largest Frobenius norm, there are three pairs with significant idiosyncratic epistasis: 124-128, 127-128, 128-129. Under the assumption that there is no link between these two quantities, it leads to a p-value equal to 0.0036 (Material and method). Therefore, it seems that the most interacting pairs of sites predicted by the Potts model within the α-helix correspond to the pairs of sites where local idiosyncratic interactions seem to result in the long term in some specific coevolution patterns between pairs of sites, which are captured by Potts model. However, these effects are not captured at the scale of the interactions between two specific sites and two specific amino acids, but at the scale of the sites.

## Discussion

The deep mutational scan we have performed here to study mutation effects in a local alpha-helix of the beta-lactamase TEM-1 reveals that epistasis is pervasive. We found that once we exclude mutations carrying irrevocable loss of function, 83% of mutations showed some strong signature of epistasis. Interestingly, though we work on a small fraction of the protein, most epistasis do not result from idiosyncratic interactions between sites, but are mostly captured by a global model of epistasis. In that model, the phenotypic impact of mutant adds up in double mutants but the non-linear translation of phenotype to fitness results in epistasis [ Wylie and Shakhnovich, 2011, Otwinowski, McCandlish and Plotkin, 2018]. The functional form of the non-linear mapping between the fitness and the phenotype may reflect the global impact of the mutations on the protein stability, in particular for the secondary structure component under investigation, and on its functionality. The phenotype to fitness mapping therefore reflects the environmental pressure on the activity of the protein, tuned by the experimental conditions, here determined by the antibiotic concentration [Otwinowski, McCandlish and Plotkin, 2018, Stiffler, Hekstra and Ranganathan 2015, Roussel *et al* 2021]. Using the two-state model and the single and double mutations scan, we could estimate for each single mutant a phenotypic effect in the form of an energy change, ΔΔG. Within this model, we could explain both qualitatively and quantitatively a large fraction of the observed epistasis (rho=0.81).

Moreover, as, according to the two-state model, the mutational effects on the phenotype are additive, we could fit the ΔΔG parameters only from the single mutational data, to predict epistasis with a good accuracy as estimated with a Spearman correlation ranging from 0.6 to 0.7.

The large contribution of this global epistasis we observed despite our focus on a local structure of the protein is remarkable and further emphasizes the importance of this form of epistasis, whose overall relative contribution should only increase as we consider larger fractions of the protein. The importance of these macroscopic form of epistasis at the protein level is reminiscent of the negative epistasis found genome-wide in experimental evolution (Chou *et al*., 2011; Khan *et al*., 2011; Wiser, Ribeck and Lenski, 2013; Kryazhimskiy *et al*., 2014).

Our precise estimates of log-fitness allowed us to identify some deviations to the two-state model. Interestingly, there was also some consistency in these deviations that were more likely to occur between residues in direct contacts in the protein structure. We found for instance some examples of local interactions linked to charge conservation. Deviation from the additivity at the phenotypic level may generate these deviations from macroscopic epistasis. We would like to point out that our alpha helix is not included in the active site of the protein. We believe that our two-state model would be less predictive for sites included in the active site, where activity would predominate over global epistasis (Rodrigues *et al*., 2016). However, we estimate that for a majority of sites, this global epistasis dominates.

Both global epistasis and deviation from it seem to be connected to the 3D structure of the alpha-helix under investigation either through the impact of mutations on protein stability or through contacts between the residues. Because such structure is highly conserved, we then questioned whether the determinants of epistasis were conserved enough to be detected from the analysis of Multiple Sequence Alignments (MSA) of distant homologues that share the same fold. Interestingly, both the signature of the macroscopic model and the patterns of deviations were recovered through the integration of MSA in the Potts model. First, the estimated ΔΔG correlated linearly with the Potts model mutation energy predictions. However, because macroscopic epistasis results from a precise non-linear mapping of phenotype to fitness, the Potts model estimates of ΔΔG had to be inserted in the two-state model to have some predictive power on the observed epistasis (mostly on the sign of epistasis). Second, pairs of sites that showed the strongest signal of coevolution through evolutionary times (as measured through the Frobenius norm of the couplings of Potts model) were the ones that deviated the most from the macroscopic model. These idiosyncratic epistatic interactions seem therefore to generate in the long-term some co-evolution patterns between pairs of sites that can be captured by models trained on MSA.

The fact that the experimental epistasis we characterized as either global or idiosyncratic can both be recovered to some extent from the analysis of distant homologes is telling that the molecular determinants of epistasis are long-lasting. It suggests that the persistence of the underlying mechanistic selective pressures has been long and strong enough to shape the long-term evolution of the protein family. Despite the wide-spread level of epistasis we recovered in our data, these observations reject a model in which epistatic interactions are fully volatile and change quickly with protein sequence as suggested for instance in the NK model(Kauffman and Weinberger, 1989). Our data suggest a rather smooth and consistent protein mutational landscape. This offers the hope that its property could be tractable and extrapolated from one homologue to another using combinations of mutational scans and in-depth multiple sequence alignment analysis (Cocco, Posani and Monasson 2021).

## Material and Methods

### Construction of a library of barcoded mutants

The sequence of TEM-1 was mutagenized using a previously published phagemid (Firnberg and Ostermeier, 2012) that was slightly modified. This phagemid allows high throughput mutagenesis to be performed, from a phage mediated single stranded amplification, and the synthesis of the other strand using a pool of mutated oligonucleotides. For our purposes, these oligonucleotides each carry two degenerate NNS codons (N is either A, T, G or C; S is either G or C) in the alpha helix of interest. A collection of 150 000 mutants was made with this protocol. The mutants were then combined through Gibson assembly to a genetic barcode of sequence NNNNNATNNNNNATNNNNNATNNNNN flanking a gene providing resistance to the antibiotic chloramphenicol. Two million barcoded mutants were recovered (see supplement for more details).

### Coupling barcode sequences to mutant sequence

To find for each barcode the mutations in the alpha helix it is linked to, two independent PCR (including one in emulsion) were done with one end of the product corresponding to the barcode and the other end to the alpha helix sequence. Using paired end sequencing with Miseq technology, both barcode sequence and alpha helix sequence of the mutants could be recovered. Each barcode was associated with an alpha helix sequence. To prevent wrong association of some barcodes to diverse alpha helix sequences due to recombination that may occur during the PCR, we excluded from the analysis barcodes for which the second most frequent alpha helix sequence had a frequency higher than 20% (see supplement for more details).

### Selection experiment

To infer fitness of these mutants, a competition experiment was performed. The collection was grown in 100ml of MH broth in flasks to an Optical Density (OD) of 0.4 in the absence of amoxicillin, and subsequently diluted 32-fold in 100ml of MH broth this time supplemented with 8 g/l amoxicillin. The optical density was followed through time and as soon as OD reached 0.2, a 32-fold dilution was performed in fresh media with the same concentration of antibiotics. Up to 6 cycles were performed, corresponding to about 30 generations. At each dilution, samples were taken to purify the plasmid and sequence the barcode. Barcodes’ sequences were then clustered and transformed into counts for their respective alpha helix sequences. For a given alpha helix sequence, multiple barcodes could be used to estimate fitness through change in frequency through time. A simple scan of the first cycle of selection was used to eliminate, for each alpha helix mutant sequence, the barcodes that clearly deviated from the pattern of frequency changes observed for that mutant (see supplement for more details).

### Inference of log-fitness

For a given mutant i, we consider that the total population of plasmids *N*_*i*_ carrying this mutant follows an exponential growth

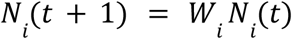

where *W*_*i*_ denotes the absolute fitness of the mutant i. However, we do not have access to the total population over time but to some measurements of the population 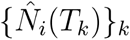 at different times *T*_*k*_ sampled by a DNA sequencer. Consequently, we construct an inference procedure to estimate its absolute fitness *W*_*i*_ knowing the measurements of the population 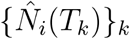 at different times *T*_*k*_. The probability of 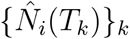 at different times *T*_*k*_ knowing *W*_*i*_ can be written (see Supplementary “Inference procedure of the log-fitness”), as a binomial distribution:

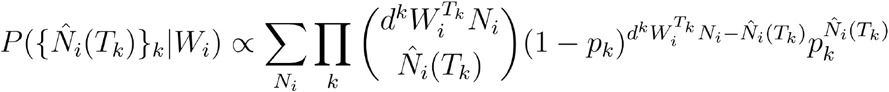

where d denotes a dilution ratio and *p*_*k*_ the sampling rate of the DNA sequencer at time *T*_*k*_.

The absolute fitness are estimated by maximizing the following likelihood

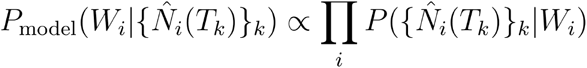

### Inference of ΔΔ*G* on single and double mutants

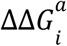 and Δ*G*_0_ are estimated by minimizing the following cost function, which corresponds to a robust nonlinear regression

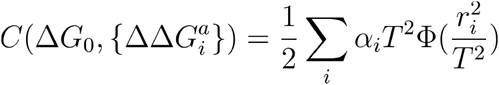

where *r*_*i*_ is the residue

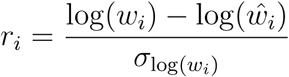

with log(w) the log-fitness of the mutant, log(\hat w) the theoretical log-fitness of the mutant given by the two-state model (Equation 3) and \sigma_log(w) the standard deviation of the log-fitness inferred with our inference procedure.

If α_*i*_ = 1 and Φ(*x*) = *x*, the cost function corresponds to the canonical least-squares estimation. However, in our data, we have strong outliers, and as we perform the inference on single and double mutants, single mutants are underrepresented compared to double mutants, which are more numerous.

To penalize the strong outliers, we used Φ(*x*) = arctan (as a loss. The parameter T is a threshold that controls the importance of the regularization of the outliers and is chosen such that 30% of the mutations are considered as outliers. The results are consistent for a wide range of thresholds T (from T = 20 to T = 100), penalizing only the strong outliers. For the parameters shown in the paper, T=50.

In order to give the same importance to single and double mutants, we used a statistical weight α_*i*_. For the double mutants, α_*i*_ = 2. For the single mutants, α_*i*_ is equal to the number of double mutants with this single mutation.

### Selection of mutants for ΔΔ*G* and Δ*G*_0_ inference

For the inference of the ΔΔ*G* and Δ*G*_0_ from the single mutational scan we kept all single mutants with a log-fitness greater than -0.6. For the inference of the same parameters from the single and double mutants, we add to the fit all the double mutations for which the single mutants are kept; but, if their log-fitness is smaller than -0.6 we threshold it to -0.6.

### Selection of mutants for epistasis

To have a high resolution on epistasis, we calculate it only for double mutants that have log-fitness greater than -0.6 and whose two associated single mutants have log-fitness greater than -0.6.

### Inference of independent model and Potts model

All models are trained by maximizing the log-likelihood of MSA built on homologs of class A β-lactamases cleaned by hand (Philippon *et al*., 2016, 2019) enriched on SwissProt and TrEMBL (The UniProt Consortium, 2021), with a total of B = 8749 sequences with length L = 253. Each sequence is reweighted according to the classical reweighting scheme ((Morcos *et al*., 2011), with a threshold equals to 0.2, leading to an effective number of sequences B_eff = 2480. For Potts model, the log-likelihood was maximized with Pseudolikelihood maximization (Ekeberg *et al*., 2013; Ekeberg, Hartonen and Aurell, 2014) with *L*_2_ regularization (for the couplings, 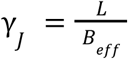, for the fields, 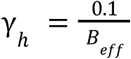) and color compression (Rizzato *et al*., 2020), with a threshold *f*_0_ = 0.

## Supporting information

Supplementary material: details of experiments

Supplementary material: details of log-fitness inference

## Supplementary figures

**Supplementary Figure A:**
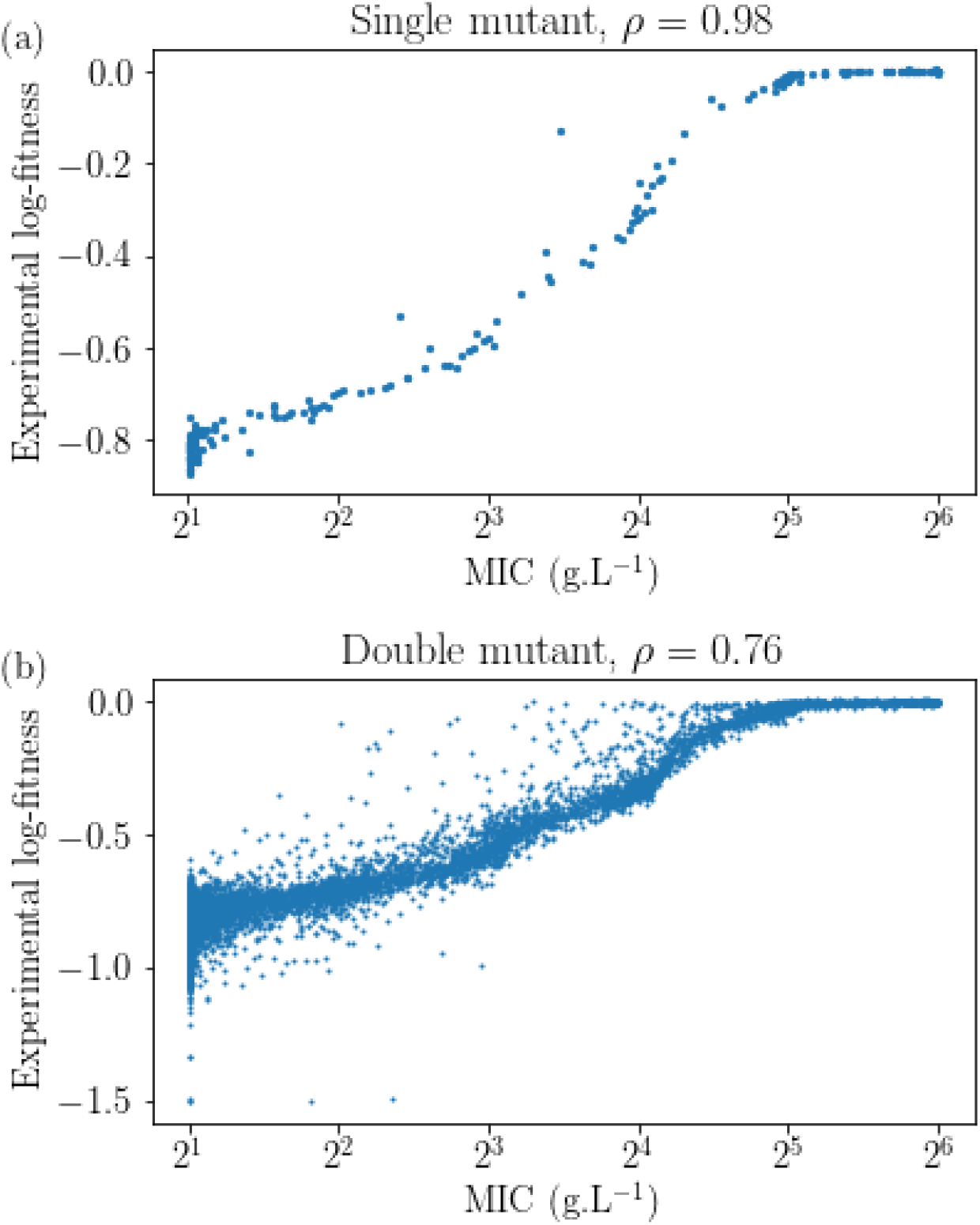
Comparison between experimental log-fitness and MIC. (a) For single mutants. (b) For double mutants.

**Supplementary Figure B:**
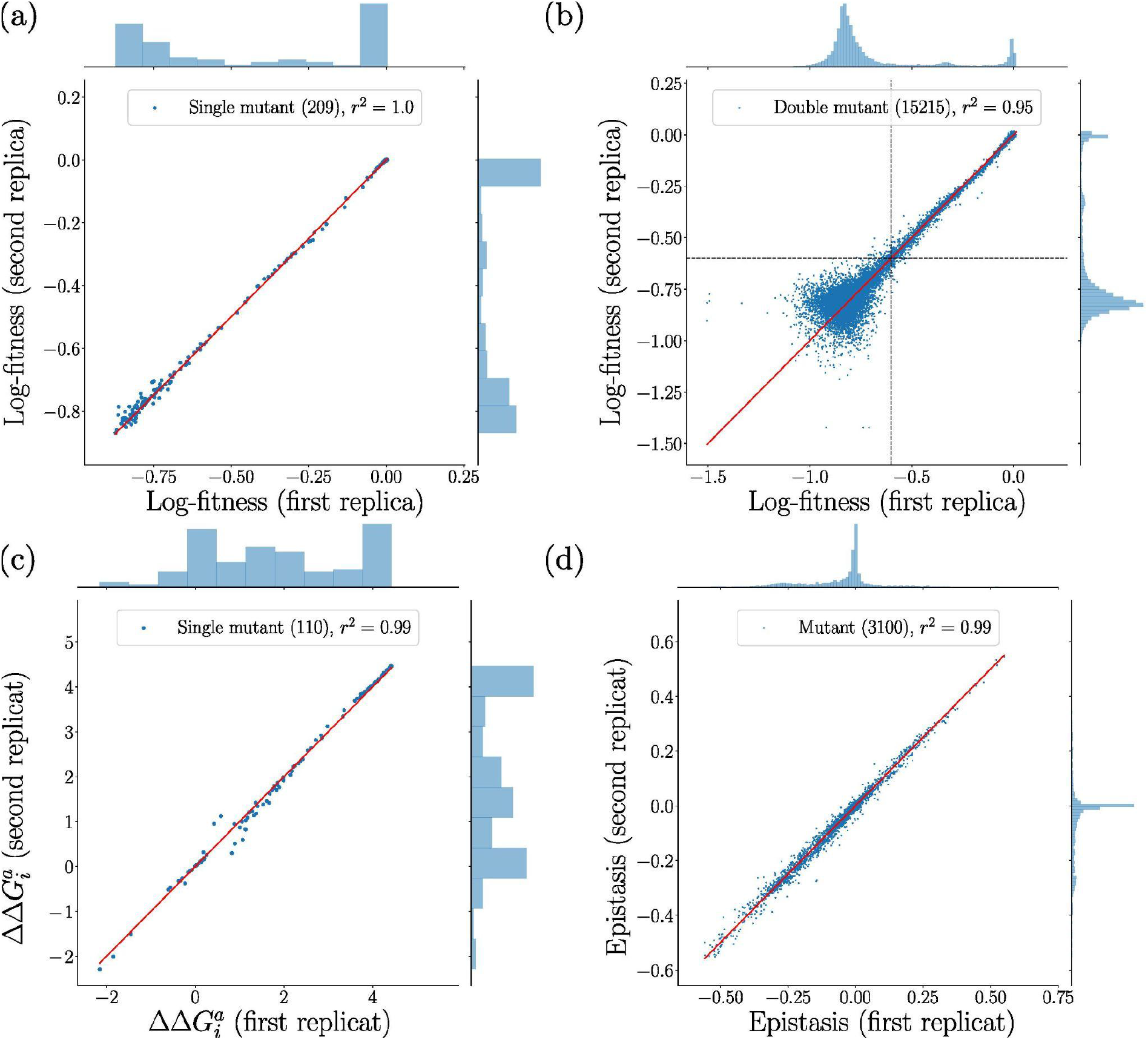
Comparison between two biological semi-replicates. (a) Single mutants’ log-fitness. (b) Double mutants’ log-fitness. (c) ΔΔ*G*. (d) Epistasis.

**Supplementary Figure C:**
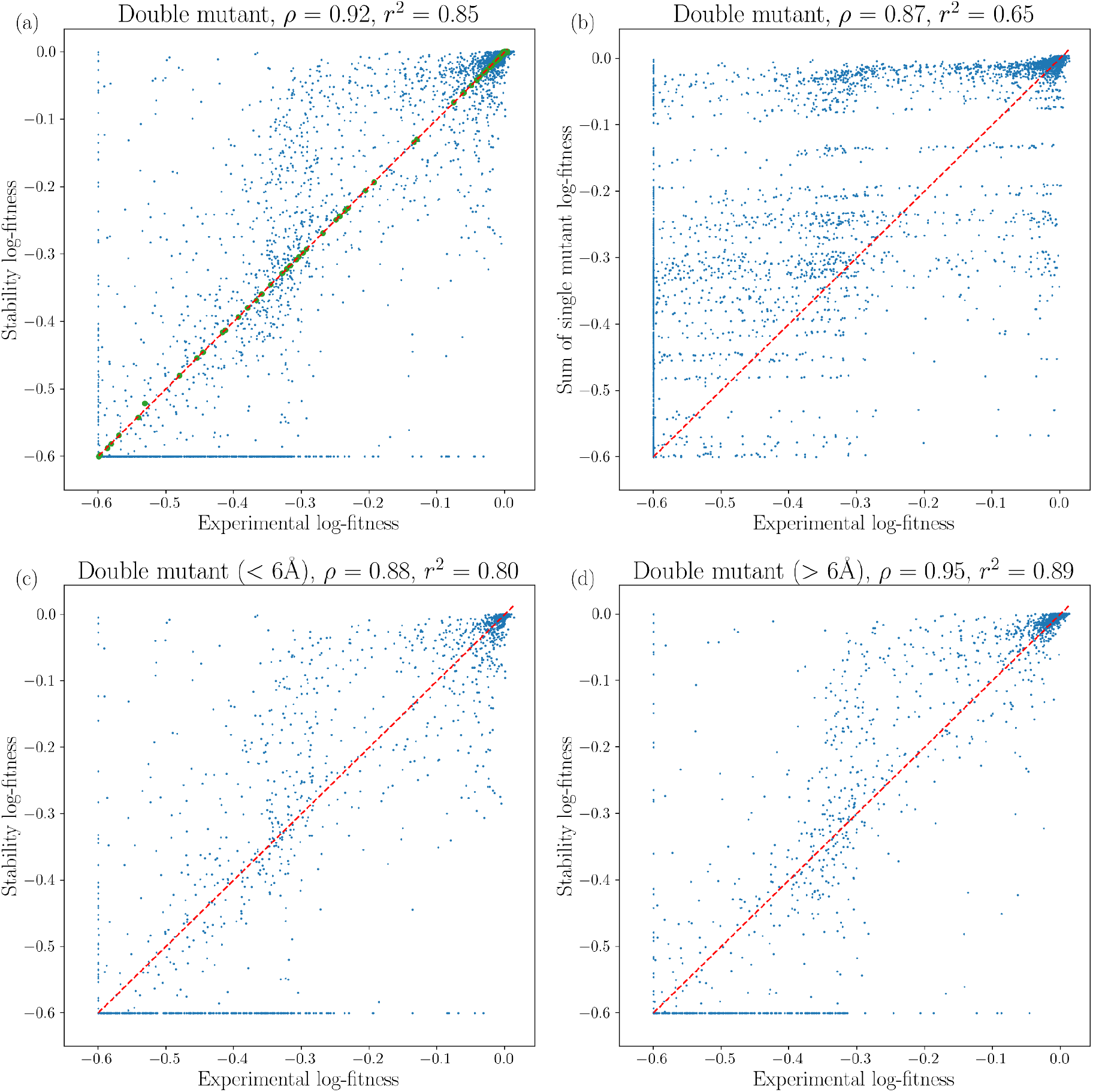
Comparison between experimental log-fitness and stability model. Green dots, single mutant. Blue dots, double mutant. Red dashed line, identity. (a) Stability log-fitness versus experimental log-fitness. (b) Sum of single mutant log-fitness versus experimental log-fitness. (c) Stability log-fitness versus experimental log-fitness, close pairs (>6Å). (d) Stability log-fitness versus experimental log-fitness, distant pairs (>6Å).

**Supplementary Figure D:**
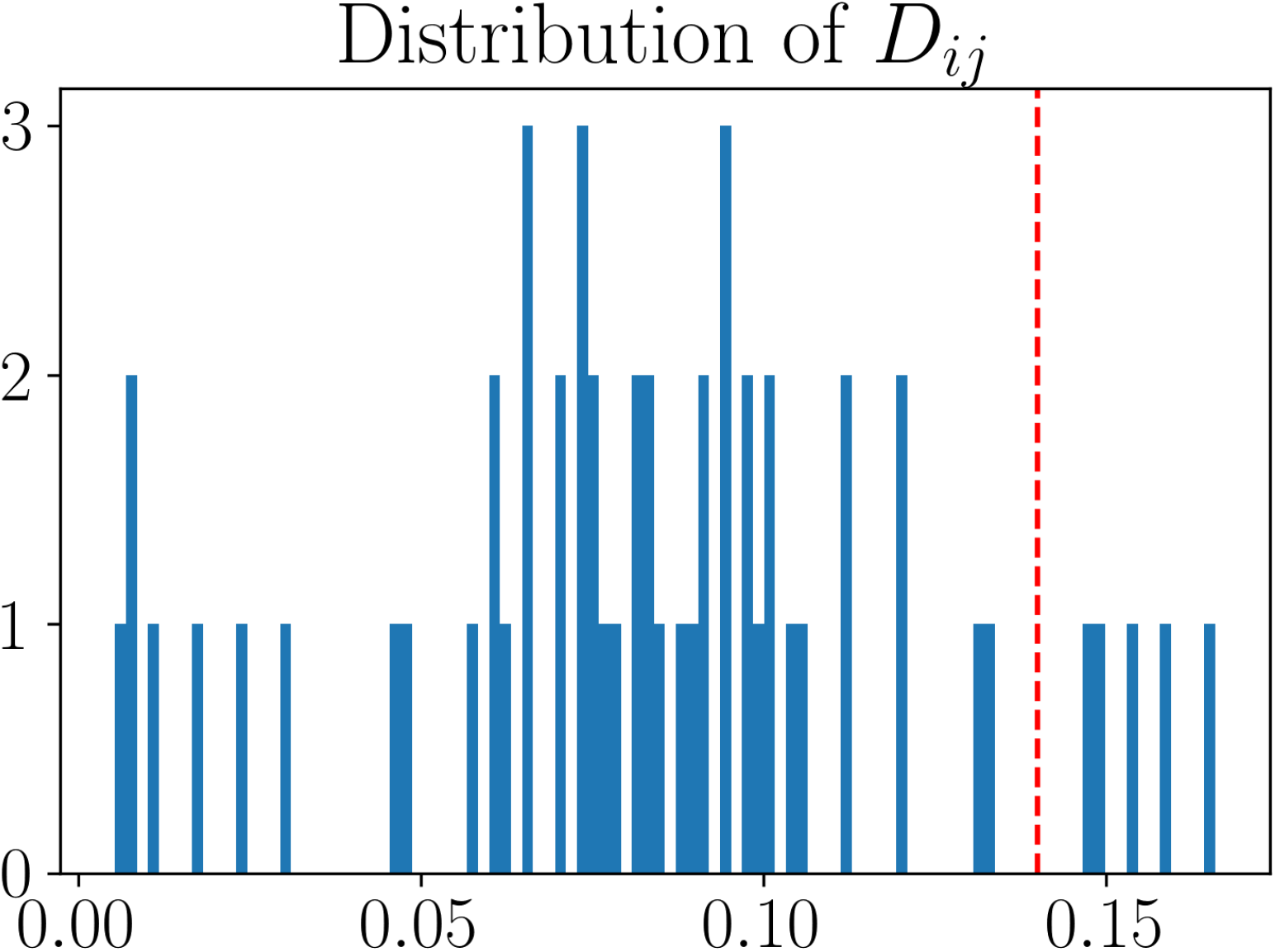
Distribution of *D*_*ij*_. Red dashed line separates the top five pairs of sites from the others.

**Supplementary Figure E:**
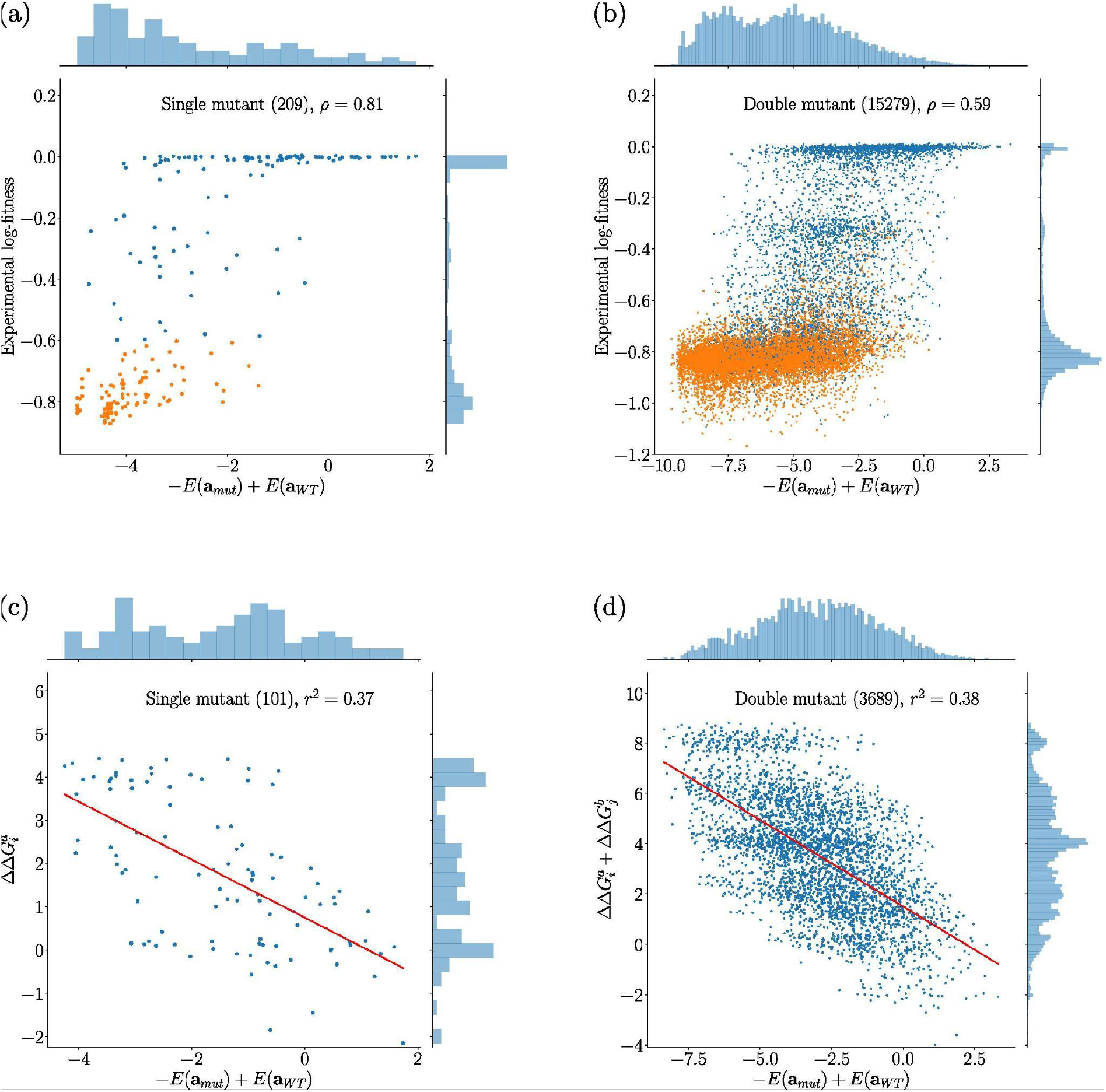
Independent model’s energies versus experimental quantities Blue dots correspond to mutations we used to estimateΔΔ*G*. Orange points are the other experimental mutations. For panels (c) and (d), only mutations present in the MSA are kept (with a different background than TEM-1), decreasing the number of single mutants from 111 to 101 and the number of double mutants from 4392 to 3689. (a) Experimental log-fitness against 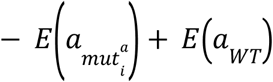 for single mutants. (b) Experimental log-fitness against 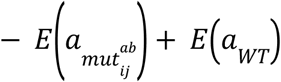 for double mutants. (c) 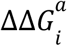 against 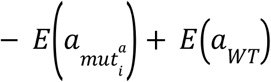 (d) 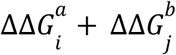 against 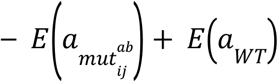.

**Supplementary Figure F:**
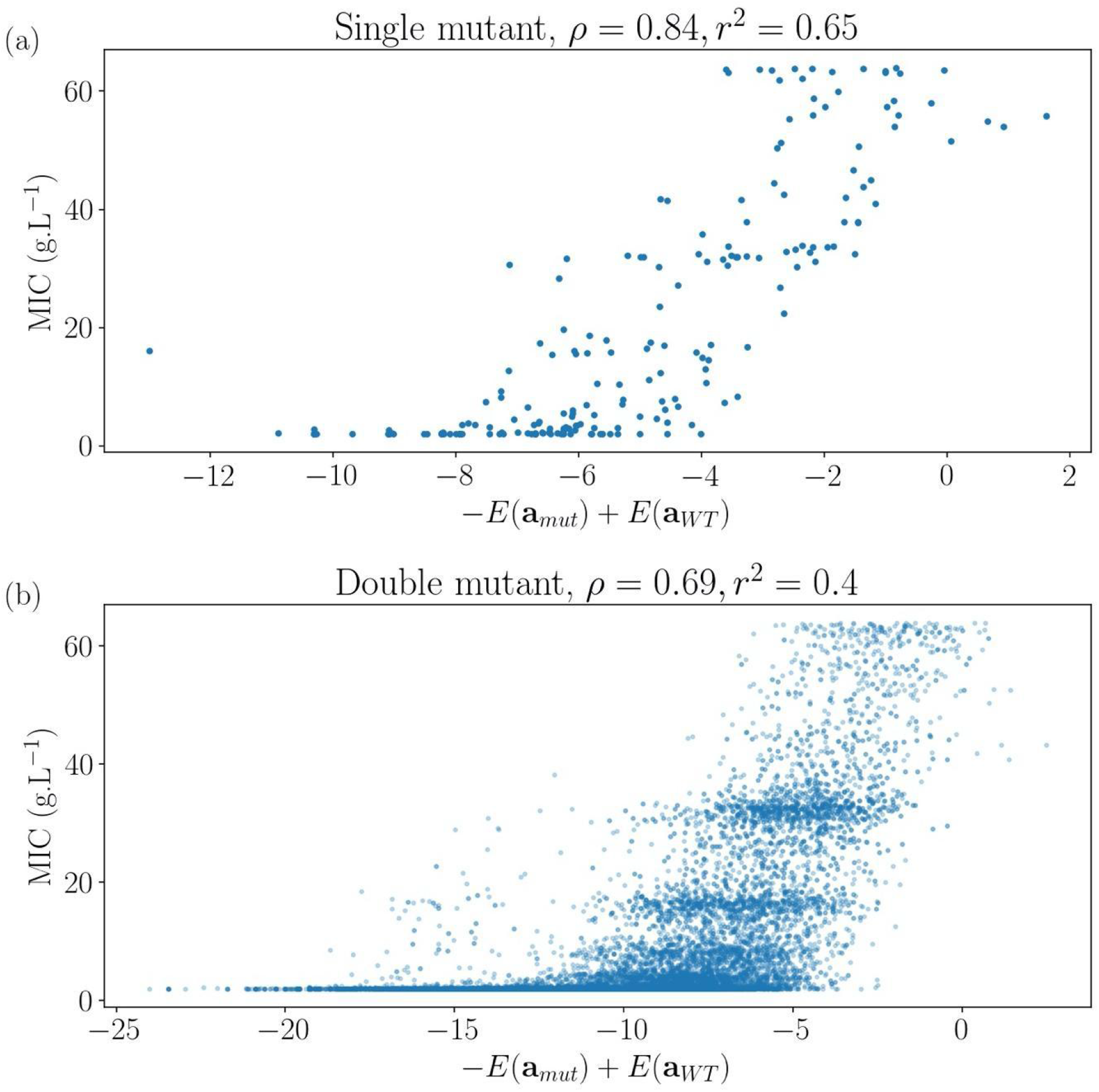
Comparison between MIC and Potts’ energies. (a) For single mutants. (b) For double mutants.

## Bibliography

Bank, C. et al. (2015) ‘A Systematic Survey of an Intragenic Epistatic Landscape’, Molecular Biology and Evolution, 32(1), pp. 229–238. doi:10.1093/molbev/msu301.

Bank, C. et al. (2016) ‘On the (un)predictability of a large intragenic fitness landscape’, Proceedings of the National Academy of Sciences, 113(49), pp. 14085–14090. doi:10.1073/pnas.1612676113.

Bershtein, S., Goldin, K. and Tawfik, D.S. (2008) ‘Intense neutral drifts yield robust and evolvable consensus proteins’, Journal of Molecular Biology, 379(5), pp. 1029–1044. doi:10.1016/j.jmb.2008.04.024.

Blanquart, F. et al. (2014) ‘Properties of selected mutations and genotypic landscapes under Fisher’s geometric model’, Evolution, 68(12), pp. 3537–3554. doi:10.1111/evo.12545.

Bloom, J.D. et al. (2005) ‘Thermodynamic prediction of protein neutrality’, Proceedings of the National Academy of Sciences of the United States of America, 102(3), pp. 606–611. doi:10.1073/pnas.0406744102.

Chou, H.-H. et al. (2011) ‘Diminishing returns epistasis among beneficial mutations decelerates adaptation’, Science (New York, N.Y.), 332(6034), pp. 1190–1192. doi:10.1126/science.1203799.

Cocco S., Posani L., Monasson R. ‘Functional couplings from sequence and mutational data.’ In preparation (2021)

DePristo, M.A., Weinreich, D.M. and Hartl, D.L. (2005) ‘Missense meanderings in sequence space: a biophysical view of protein evolution’, Nature Reviews Genetics, 6(9), pp. 678–687. doi:10.1038/nrg1672.

Dunn, S.D., Wahl, L.M. and Gloor, G.B. (2008) ‘Mutual information without the influence of phylogeny or entropy dramatically improves residue contact prediction’, Bioinformatics, 24(3), pp. 333–340. doi:10.1093/bioinformatics/btm604.

Ekeberg, M. et al. (2013) ‘Improved contact prediction in proteins: Using pseudolikelihoods to infer Potts models’, Physical Review E, 87(1), p. 012707. doi:10.1103/PhysRevE.87.012707.

Ekeberg, M., Hartonen, T. and Aurell, E. (2014) ‘Fast pseudolikelihood maximization for direct-coupling analysis of protein structure from many homologous amino-acid sequences’, Journal of Computational Physics, 276, pp. 341–356. doi:10.1016/j.jcp.2014.07.024.

Figliuzzi, M. et al. (2016) ‘Coevolutionary Landscape Inference and the Context-Dependence of Mutations in Beta-Lactamase TEM-1’, Molecular Biology and Evolution, 33(1), pp. 268–280. doi:10.1093/molbev/msv211.

Firnberg, E. and Ostermeier, M. (2012) ‘PFunkel: efficient, expansive, user-defined mutagenesis’, PloS One, 7(12), p. e52031. doi:10.1371/journal.pone.0052031.

Fowler, D.M. and Fields, S. (2014) ‘Deep mutational scanning: a new style of protein science’, Nature Methods, 11(8), pp. 801–807. doi:10.1038/nmeth.3027.

Gros, P.-A., Le Nagard, H. and Tenaillon, O. (2009) ‘The evolution of epistasis and its links with genetic robustness, complexity and drift in a phenotypic model of adaptation’, Genetics, 182(1), pp. 277–293. doi:10.1534/genetics.108.099127.

Hagen, J.B. (2000) ‘The origins of bioinformatics’, Nature Reviews. Genetics, 1(3), pp. 231–236. doi:10.1038/35042090.

von Heijne, G. (1991) ‘Proline kinks in transmembrane α-helices’, Journal of Molecular Biology, 218(3), pp. 499–503. doi:10.1016/0022-2836(91)90695-3.

Hopf, T.A. et al. (2017) ‘Mutation effects predicted from sequence co-variation’, Nature Biotechnology, 35(2), pp. 128–135. doi:10.1038/nbt.3769.

Jacquier, H. et al. (2013) ‘Capturing the mutational landscape of the beta-lactamase TEM-1’, Proceedings of the National Academy of Sciences of the United States of America, 110(32), pp. 13067–13072. doi:10.1073/pnas.1215206110.

Kauffman, S.A. and Weinberger, E.D. (1989) ‘The NK model of rugged fitness landscapes and its application to maturation of the immune response’, Journal of Theoretical Biology, 141(2), pp. 211–245. doi:10.1016/s0022-5193(89)80019-0.

Khan, A.I. et al. (2011) ‘Negative epistasis between beneficial mutations in an evolving bacterial population’, Science (New York, N.Y.), 332(6034), pp. 1193–1196. doi:10.1126/science.1203801.

Kryazhimskiy, S. et al. (2014) ‘Microbial evolution. Global epistasis makes adaptation predictable despite sequence-level stochasticity’, Science (New York, N.Y.), 344(6191), pp. 1519–1522. doi:10.1126/science.1250939.

Martin, G., Elena, S.F. and Lenormand, T. (2007) ‘Distributions of epistasis in microbes fit predictions from a fitness landscape model’, Nature Genetics, 39(4), pp. 555–560. doi:10.1038/ng1998.

Morcos, F. et al. (2011) ‘Direct-coupling analysis of residue coevolution captures native contacts across many protein families’, Proceedings of the National Academy of Sciences, 108(49), pp. E1293–E1301. doi:10.1073/pnas.1111471108.

Olson, C. A., Wu, N. C. & Sun, R. A comprehensive biophysical description of pairwise epistasis throughout an entire protein domain. Curr. Biol. 24, 2643–2651 (2014).

Otwinowski, J., McCandlish, D.M. and Plotkin, J.B. (2018) ‘Inferring the shape of global epistasis’, Proceedings of the National Academy of Sciences of the United States of America, 115(32), pp. E7550–E7558. doi:10.1073/pnas.1804015115.

Philippon, A. et al. (2016) ‘A Structure-Based Classification of Class A beta-Lactamases, a Broadly Diverse Family of Enzymes’, Clinical Microbiology Reviews, 29(1), pp. 29–57. doi:10.1128/CMR.00019-15.

Philippon, A. et al. (2019) ‘Structure-based classification of class A beta-lactamases, an update’, Current Research in Translational Medicine, 67(4), pp. 115–122. doi:10.1016/j.retram.2019.05.003.

Privalov, P.L. (1979) ‘Stability of Proteins Small Globular Proteins’, in Anfinsen, C.B., Edsall, J.T., and Richards, F.M. (eds) Advances in Protein Chemistry. Academic Press, pp. 167–241. doi:10.1016/S0065-3233(08)60460-X.

Rizzato, F. et al. (2020) ‘Inference of compressed Potts graphical models’, Physical Review E, 101(1), p. 012309. doi:10.1103/PhysRevE.101.012309.

Rodrigues, J. V. et al. Biophysical principles predict fitness landscapes of drug resistance. Proc. Natl Acad. Sci. USA 113, E1470–E1478 (2016).

Roussel et al. (2021), Epistasis at different selection pressures in an α-helix of β-lactamase TEM-1 ‘ in preparation.

Salinas, V.H. and Ranganathan, R. (2018) ‘Coevolution-based inference of amino acid interactions underlying protein function’, eLife. Edited by N. Ben-Tal and D. Weigel, 7, p. e34300. doi:10.7554/eLife.34300.

Salverda, M.L.M., De Visser, J.A.G.M. and Barlow, M. (2010) ‘Natural evolution of TEM-1 β-lactamase: experimental reconstruction and clinical relevance’, FEMS microbiology reviews, 34(6), pp. 1015–1036. doi:10.1111/j.1574-6976.2010.00222.x.

Sarkisyan, K.S. et al. (2016) ‘Local fitness landscape of the green fluorescent protein’, Nature, 533(7603), pp. 397–401. doi:10.1038/nature17995.(Sarkisyan et al., 2016)

Stiffler, M. A, Hekstra D.R., Ranganathan R. Evolvability as a function of purifying selection in TEM-1 β-lactamase, Cell 160 (5):882–892 (2015)

Tenaillon, O. (2014) ‘The Utility of Fisher’s Geometric Model in Evolutionary Genetics’, Annual Review of Ecology, Evolution, and Systematics, 45(1), pp. 179–201. doi:10.1146/annurev-ecolsys-120213-091846.

The UniProt Consortium (2021) ‘UniProt: the universal protein knowledgebase in 2021’, Nucleic Acids Research, 49(D1), pp. D480–D489. doi:10.1093/nar/gkaa1100.

de Visser, J.A.G.M. and Elena, S.F. (2007) ‘The evolution of sex: empirical insights into the roles of epistasis and drift’, Nature Reviews. Genetics, 8(2), pp. 139–149. doi:10.1038/nrg1985.

de Visser, J.A.G.M. and Krug, J. (2014) ‘Empirical fitness landscapes and the predictability of evolution’, Nature Reviews. Genetics, 15(7), pp. 480–490. doi:10.1038/nrg3744.

Wiser, M.J., Ribeck, N. and Lenski, R.E. (2013) ‘Long-term dynamics of adaptation in asexual populations’, Science (New York, N.Y.), 342(6164), pp. 1364–1367. doi:10.1126/science.1243357.

Wylie, C.S. and Shakhnovich, E.I. (2011) ‘A biophysical protein folding model accounts for most mutational fitness effects in viruses’, Proceedings of the National Academy of Sciences of the United States of America, 108(24), pp. 9916–9921. doi:10.1073/pnas.1017572108.

Zhao, V.Y. et al. (2021) ‘Switching an active site helix in dihydrofolate reductase reveals limits to sub-domain modularity’, Biophysical Journal [Preprint]. doi:https://doi.org/10.1016/j.bpj.2021.09.032.

