## Supplementary material: details of experiments for "Origins and breadth of pairwise epistasis in an α-helix of β-lactamase TEM-1"

#### Strains and plasmids

Strains: *E. coli* strains used in this study: **XL1-Blue** (Agilent, Santa Clara, CA) of genotype recA1 endA1 gyrA96 thi-1 hsdR17 supE44 relA1 lac [F’ proAB lacI1qZΔM15 Tn10 (Tetr)]; **CJ236** (new England biolabs) of genotype FΔ(HindIII)::cat (Tra+ Pil+ CamR)/ ung-1 relA1 dut-1 thi-1 spoT1 mcrA; **DH5α** (Invitrogen) of genotype F– Φ80lacZΔM15 Δ(lacZYA-argF) U169 recA1 endA1 hsdR17 (rK–, mK+) phoA supE44 λ– thi-1 gyrA96 relA1; **Dh10b** Electromax (ThermoFisher Scientific) of genotype $F^-$*mcr*A Δ(*mrr*-*hsd*RMS-*mcr*BC) Φ80*lac*ZΔM15 Δ*lac*X74 *rec*A1 *end*A1 *ara*D139Δ(*ara*, *leu*)7697 *gal*U *gal*K $\lambda^-$*rps*L *nup*G

Plasmids : Phagemid **pSkunk3-TEM-1** was obtained graciously from Elad Firnberg and Marc Ostermeier. The **pSkunk-TEM-helix** was created by inserting a *Nco*I restriction site 2 bases before the beginning of alpha helix to mutagenize using single-step Pfunkel mutagenesis. Using the same protocol, we also inserted *Xho*I and *Not*I restrictions sites which surround the streptomycin/spectinomycin (Sm/Spec) resistance gene. Plasmid pKD3 was used to amplify the cat gene encoding chloramphenicol (Cm) resistance. The **pSkunk-TEM-helix-Cm** was created by swapping the Sm/Spec resistance gene with a Cm resistance gene and adding a DNA barcode of 20 degenerate nucleotides.

#### Mutagenesis

To produce the mutants, we used the PFunkel mutagenesis strategy (Datsenko and Wanner, 2000**). ssDNA production:** Uracil-containing single-strand DNA of pSkunk3-TEM-helix was produced as published by Firnberg and Ostermeier, 2012; Kowalsky *et al.*, 2015 (except the final centrifugation step, which was performed at 26200g for 1h at 4°C). DNA was quantified using the Qubit® ssDNA Assay Kit (ThermoFisher Scientific).

**Single step Pfunkel mutagenesis:** Mutagenesis was performed as previously described with 1 µg of ssDNA used as template in a total volume of 100 µl. The only difference was the elongation step which was 15 min. Thus, the reaction cycling conditions were 95°C for 3 min, followed by 55°C for 90 sec, and 68°C for 15 min and 45°C for 15 min (Kowalsky *et al.*, 2015). We used the innuPREP PCRpure Kit (analytik jena) to purify DNA and eluted in 15 µl of distilled DNAse/RNAse free water. 2 µl were then electroporated in 20 µl of DH5α electrocompetent cells and then incubated with 500 µl of LB media for 1 hour at 37°C with shaking at 250 rpm. The transformation was plated on LB agar with 50 µg/ml streptomycin and then incubated overnight at 37°C. To check for mutagenesis efficiency, PCR verification and Sanger sequencing was performed on isolated colonies. Mutants were stocked in LB-glycerol 40% after an overnight culture at 37°C in LB media containing 50 µg/ml streptomycin.

**Comprehensive pfunkel mutagenesis:** Primers containing all combinations of two NNS degenerate codons (N is either A, T, G or C; S is either G or C) of the 11 codons of mutagenized alpha helix were designed with 20 fixed bp surrounding the alpha helix (on 5' and 3').

Protocols were performed as published by Firnberg and Ostermeier, 2012; Kowalsky *et al.*, 2015 for single site mutagenesis with amplification step of 20 min. Purification was carried out using the innuPREP PCRpure Kit (analytik jena) and DNA was eluted in 15µl of distilled water. 2 µl were electroporated in 20 µl of Dh5a electrocompetent cells and then incubated with 500 µl of LB media for 1 hour at 37°C with shaking at 250 rpm. The transformation was plated on LB agar with 50 µg/ml streptomycin and then incubated overnight at 37°C. A pool of 150 000 colonies was scraped from the LB agar plates (245 mm X 245 mm, Greiner bio-one) in LB broth and frozen at -80°C in LB/glycerol 40%. After pooling all colonies together, plasmids were extracted from an aliquot using plasmid miniprep (Qiagen, Valencia, CA), forming the library of mutants.

**Mutant barcoding:** 10 µg of plasmid extraction of the library of mutants was digested with *Not*I, *Xho*I and *Nco*I (buffer 3.1), in 500ul total reaction volume (New England Biolabs), gel extracted (band of 3350 bp) using Qiagen Gel Extraction kit and then also cleaned a 2nd time with Qiagen PCR Purification kit. The final concentration was : 25 ng/ul.

pKD3 was used as template for PCR amplification of Cat using specific primers that also contained overlapping regions of pSkunk-TEM-helix (for subsequent Gibson Assembly). The forward primer also contained a non-overlapping region with a DNA barcode consisting of 20 degenerated nucleotides. Phusion® High-Fidelity DNA Polymerase (New England Biolabs) was used with reaction cycling conditions : 98°C for 30 sec, followed by 35 cycles of 98°C for 10 sec, and 62°C for 30 sec, 72°C for 15 sec and a final extension at 72°C for 2 min.

The plasmid **pSkunk-TEM-helix-Cm** was created by switching the spectinomycin/streptomycin resistance with the Cm resistance cassette amplified previously using Gibson Assembly. This allows the integration of the DNA barcode (Gibson *et al.*, 2009)(New England Biolabs). Gibson reaction was carried out with 3µl of 25 ng/ul of plasmid fragment and 1.3 µl of 88 ng/µl of barcode-CmR-amplicon (1:5 molar ratio backbone:insert), in a total of 20 µl reaction mix and incubated at 50°C for 1 hour. The total volume of Gibson reaction was dialysed with water for 30 mins and 4 µl was electroporated in 20 µl of Dh10b Electromax competent cells that were then incubated in 500 µl of LB media for 1 hour at 37°C with shaking at 250 rpm. The transformants were plated on LB agar with 25 µg/ml chloramphenicol and then incubated overnight at 37°C. A pool of about $2.10^6$ colonies was scraped from the LB agar plates (245 mm X 245 mm, Greiner bio-one) in LB broth and frozen at -80°C in LB/glycerol 40%.

**Selection experiments**

To estimate mutant fitness, a selection experiment was performed.

**Selection with 8g/l of amoxicillin:** 1ml of the frozen barcoded library cells stock was cultured in MH broth at 37°C with shaking at 250 rpm to $OD_{600}$ 0.4 without antibiotics (named T0). Then, 3.2 ml of this first culture were used to re-inoculate 96.8 ml MH broth supplemented with 8g/l amoxicillin (corresponding to 2 dilutions under MIC of TEM-1) until $OD_{600}$ 0.2 (this point is called T1). This re-inoculation of 3.2 ml of culture of $OD_{600}$ 0.2 in fresh 96.8 ml MH broth supplemented with 8 g/l amoxicillin was repeated until approximately 30 total population-averaged generations (T6). Half of these cultures were pelleted for plasmid extraction and half were pelleted and re-suspended in LB-glycerol 40% for storing at -80°C. All this experiment was performed in duplicate.

**Library Preparation and Deep Sequencing**

Deep sequencing was used to obtain count data for each variant in the population.

The protocol is carried out in 2 steps:

**Combining barcodes and sequences**: the first step is to reveal which barcodes are linked to which mutations in the T0 library. For that, a two-step PCR method was used to amplify the corresponding part of the gene, including the alpha helix sequence on 5' part and barcode sequence on 3' extremity, and to add the illumina sequencing adapter and multiplex barcode sequences. In detail, plasmid DNA concentration was determined using qubit fluorometric quantification (ThermoFisher scientific) and normalized to 2.5 ng/µl. 12.5 ng of DNA was used for the 1st PCR using specific primers and allowed the attachment of an adaptor that is necessary for the 2nd PCR. Between specific primers and adaptors, 6 degenerate nucleotides were inserted in order to increase the diversity of DNA to facilitate MiSeq clustering (by improving crosstalk and phasing calculations). In order to decrease recombination that arises during PCR, a third PCR was made with an emulsion-PCR protocol (Micellula, following the manufacturer's guidelines). Kapa Hifi Hotstart Ready Mix PCR Kit polymerase (Kapa Biosystems) was used for amplification. The reaction cycling conditions were 95°C for 30 sec, followed by 12 cycles of 95°C for 10 sec, 55°C for 30 sec, 68°C for 30 sec and a final extension at 68°C for 5 min.

After gel purification using Qiagen gel extraction kit (Valencia, CA), DNA was quantified using qubit fluorometric quantification and DNA concentration was normalized. The 2nd PCR was performed using 5 ng of DNA using primers commercialized by Illumina in the Nextera Index Kit allowing the dual indexing. The reaction cycling conditions were the same as previously but only 11 cycles were performed using Kapa Hifi Hotstart Ready Mix PCR Kit polymerase (Kapa Biosystems). After gel purification with Qiagen gel extraction kit (Valencia, CA), quantification using qPCR kapa Hifi Hotstart (Kapa Biosystems) on a Light cycler 480 Roche was performed with reaction cycling conditions of 95°C for 5 min, followed by 35 cycles of 95°C for 30 sec and 60°C for 45 sec as specified by Kapa Biosystems.

This library, corresponding to the first time-point of evolution (T0), was diluted to 12 pM and loaded on the MiSeq with a mix of 10% PhiX DNA (PhiX Control v3, illumina). Three MiSeq V3 2x75 bp paired-end runs (Illumina technology) were performed for this part, resulting in a total of > 40M reads, for an expected ~20x coverage of barcode diversity. The paired-end reads are non-overlapping, with the alpha helix sequence on Read 1 and the barcode sequence on Read 2.

**Barcoding sequencing**: The second step consists of sequencing the barcodes alone at the different time-points (T0, T1, T2, T4 and T6). For this, a similar protocol was carried out using oligonucleotides that surround the barcode region; employing the same 2-step PCR based method and similar conditions. In this case, the 6 degenerate nucleotides inserted on either side of the barcode region during the 1$^{st}$ PCR also allowed us to remove PCR duplicates arising from the 2$^{nd}$ PCR. All libraries corresponding to the different evolution time-points were quantified using a qPCR-based method (integragen) and pooled in equal molar quantity. They were then sequenced on a HiSeq4000 with a 2x100 bp paired-end kit (Illumina technology) by Integragen society, to give overlapping reads of the barcode region. The run resulted in ~300M raw paired-end reads, and so ~27M for each of the 10 time-points/conditions (duplicate). This gives a barcode coverage of ~14x for each time-point/condition.

### Sequence analysis

**Barcode-mutant association** (MiSeq) The following steps were performed using the Mothur software package (https://www.mothur.org/, Schloss *et al.*, 2009): raw reads from all sequencing runs were pooled together and quality-filtered by size (>69 bases), number of uncalled bases (<3 Ns) and length of longest homopolymer stretch, an indicator of overall read quality (<13 bases). Alpha helix and barcode sequences were extracted from Read 1 and Read 2, respectively, after alignment to the reference sequences (Needleman global alignment). Reads for which either the alpha helix or barcode region contained insertions or did not generate a full alignment with the reference were discarded. The Mothur precluster algorithm was then used to cluster barcode sequences differing by a Hamming distance of 1, with the aim of correcting for PCR and sequencing errors (the potential barcode diversity is so high that the presence of immediately neighbouring sequences is very likely due to these errors). The algorithm uses sequence abundance to decide the “true” (majority) sequence for each cluster, and to decide where a sequence clusters if it has >1 immediate neighbor. After de-gapping and re-clustering barcode sequences to account for any alignment ambiguities resulting from small deletions, barcode clusters were used to build a dictionary assigning each “true” barcode sequence to an alpha helix sequence. Due to the high rate of PCR-derived recombination observed (caused by the long homologous region between the barcode region and alpha helix sequence, and resulting in molecules with swapped barcodes), a haplotype-based strategy was used for this step rather than one in which each nucleotide is considered independently. This is because the small number of mutations present in each mutant means that, at any particular position, the majority of molecules will possess the WT base, and so a high recombination rate can result in consensus alpha helix sequences in which mutant bases are assigned as WT. The efficiency of this strategy was ensured by the short length of the mutagenized region and high quality of the reads, meaning that most reads did not contain a single error in the regions of interest and so were not wasted. Briefly, a custom Python script was used to perform the following: for each barcode cluster (consisting of reads whose barcode sequences are identical to or the immediate neighbor of the inferred "true" barcode sequence), the paired alpha helix sequences were fetched; the number of occurrences of each resulting alpha helix sequence was tabulated; if the cluster contains more than 2 reads in total, the most abundant alpha helix sequence is >= 5x more abundant than the second-most abundant alpha helix sequence, and the most abundant alpha helix sequence contains no Ns, then the most abundant alpha helix sequence is assigned to the "true" barcode sequence for that cluster (else the cluster is discarded).

**Barcode counting** (HiSeq): The following steps were performed using the Mothur software package (https://www.mothur.org/, Schloss *et al.*, 2009): demultiplexed forward and reverse reads were joined into contigs using Mothur's make.contigs command with the default parameters, which takes into account the Phred score to assign (or not) a base when there is disagreement between forward and reverse reads. Contigs were then quality-filtered by size (<151bp, as longer contigs imply forward and reverse reads could not be properly overlapped), number of uncalled bases (no Ns) and length of longest homopolymer stretch, an indicator of overall read quality (<13 bases). To remove the majority of PCR duplicates arising from the 2$^{nd}$ PCR (made possible by the 6 degenerate nucleotides introduced on each side of the barcode during the 1$^{st}$ PCR), if a particular contig was present more than once, only one copy was kept. Barcode sequences were then extracted after aligning full contigs to the reference sequence (Needleman global alignment). Reads containing insertions or not generating a full alignment with the reference were discarded. Next, the Mothur precluster algorithm was used to cluster barcode sequences differing by a Hamming distance of 1, with the aim of correcting for PCR and sequencing errors, as described above for the barcode-mutant association. After de-gapping and re-clustering barcode sequences to account for any alignment ambiguities resulting from small deletions, the number of occurrences of each "true" barcode was tabulated across all time-points/conditions. Finally, a custom R script was used to merge the barcode-mutant dictionary generated above with the barcode counts table.

Based on previous work, in which we found no clear effect of synonymous mutations, we combined all synonymous mutations into a single allele.

**Quality control of barcodes.**

Multiple Barcodes (fig1) were associated to the different genotypes. Several processes may lead to variability in the signal provided by the different barcodes. First, though we used some correction and some emulsion PCR to try to correct that bias, some recombination may occur during the PCR between the part of the protein and the barcode and escape our detection procedures. Hence, a Barcode may appear to be associated to the focal genotype, but may indeed correspond to an alternative genotype. Even if a barcode is associated properly to its alpha-helix genotype, we have not sequenced the whole protein. Consequently, an undetected mutation may affect the protein elsewhere and result in a modified behavior of that barcode. To limit the effect of these outliers, that are often barcodes associated with loss of function or maximal fitness, we first did a screen to filter outlier barcodes.

For that purpose, we computed the change in the focal genotype to wild-type genotype frequency over the first cycle of evolution (T0 to T1), using the sum of all barcodes linked the focal genotype. $K_j = \left( \frac{\sum_i BC_{ij}^1}{Wt^1} \frac{Wt^0}{\sum_i BC_{ij}^0} \right)$, in which $BC_{ij}^1$ is the number of reads matching the j[th] barcode associated to genotype i at time 1, and $Wt^1$ the number of reads matching barcodes associated to wild type sequence. The value of K corresponds to an estimate of fitness over one cycle. Then, for each individual barcode we can compute based on $BC_{ij}^0$ the estimated number of reads expected at T1. If the barcode is following the overall trend we expect

$$K_{ij} = \left( \frac{BC_{ij}^1}{Wt^1} \frac{Wt^0}{BC_{ij}^0} \right) = K_j$$

We expect therefore $BC_{ij}^{1}$ to be distributed with a Poisson law of parameter $\frac{\sum_i BC_{ij}^{1}}{\sum_i BC_{ij}^{0}} BC_{ij}^{0}$ .

All barcodes, with a p-value lower than $10^{-5}$ were assumed to reject that model and to be the result of some of the artifacts previously mentioned. They were discarded and this selective process was rerun once to be sure to eliminate all outliers. Reads matching all the remaining barcodes were then combined to estimate fitness. Furthermore, barcodes with less than 10 counts for the combined time T0 and T1 were excluded, as well as mutants with less than 4 barcodes.

**Computing MIC.**

For MIC determination, we first used wild-type counts at different concentrations to estimate the change in frequency of the various genotypes with antibiotic concentration. We identified a subset of clones which increased in frequency over the wild type at the highest concentration. That set of clones was used as reference. We used a moment matching approach to identify the concentrations at which the mutant is eradicated by the antibiotic, mimicking the retention of the mutant with a step down function. In detail, the normalized change in ratio of counts towards the reference set was computed through time, leading for each mutant to a set of values of the form $x_{i0}$, $x_{i1}$, $x_{i2}$, $x_{i4}$, $x_{i8}$, $x_{i16}$, with $x_{i0}=1$ the initial normalized frequency and the other values reflecting the relative maintenance or loss of the mutant with increasing concentration. The variance $V_i$ and mean $M_i$ of $x_{i.}$ were used in the following formula to compute a quantitative MIC: $MIC_i = \frac{6}{1+\frac{V_i}{(M_i)^2}}$.

**Bibliography**

Datsenko, K.A. and Wanner, B.L. (2000) 'One-step inactivation of chromosomal genes in Escherichia coli K-12 using PCR products', *Proceedings of the National Academy of Sciences of the United States of America*, 97(12), pp. 6640–6645. doi:10.1073/pnas.120163297.

Firnberg, E. and Ostermeier, M. (2012) 'PFunkel: efficient, expansive, user-defined mutagenesis', *PloS One*, 7(12), p. e52031. doi:10.1371/journal.pone.0052031.

Gibson, D.G. *et al.* (2009) 'Enzymatic assembly of DNA molecules up to several hundred kilobases', *Nature Methods*, 6(5), pp. 343–345. doi:10.1038/nmeth.1318.

Kowalsky, C.A. *et al.* (2015) 'High-resolution sequence-function mapping of full-length proteins', *PloS One*, 10(3), p. e0118193. doi:10.1371/journal.pone.0118193.

Schloss, P.D. *et al.* (2009) 'Introducing mothur: Open-Source, Platform-Independent, Community-Supported Software for Describing and Comparing Microbial Communities', *Applied and Environmental Microbiology*, 75(23), pp. 7537–7541. doi:10.1128/AEM.01541-09.
