## Supplementary material: details of log-fitness inference for "Origins and breadth of pairwise epistasis in an α-helix of β-lactamase TEM-1"

### Supplementary material: inference
### Origins and breadth of pairwise epistasis in an $\alpha$-helix of $\beta$-lactamase TEM-1

#### 1 Inference procedure of the log-fitness

In this section, we detail the inference procedure we developed to estimate the log-fitness of each mutant. We denote as $\hat{N}_i(T_k)$ the number of plasmids measured at time $T_k$ carrying the mutation $i$ (the different barcodes encoding for one given mutation are grouped together).

##### 1.1 Modelization of the DNA sequencer

To infer the fitness of each mutation $i$, we modeled the evolution of the number of plasmids over time. $N_i(T_k)$ denotes the total population of plasmids carrying the mutation $i$ at time $T_k$

According to the definition of the absolute fitness, $N_i(T_k)$ follows an exponential growth

$$N_i(t+1) = W_i N_i(t) \ , \tag{1}$$

The relative log-fitness with respect to the wild-type reads

$$\log(w_i) = \log(W_i) - \log(W_{WT}) \ , \tag{2}$$

However, we do not have access directly to $N_i(T_k)$, the true population, but to $\hat{N}_i(T_k)$, the result of the measurement by the DNA sequencer.

According to the experimental protocol, at each time $T_k$, due to the different dilutions after each measure, the population of plasmids in the DNA sequencer at time $T_k$ can be written as $d^k W_i^{T_k} N_i$, where $d$ is the dilution factor ($d = \frac{1}{32}$).

The DNA sequencer does not sample all the plasmids, but samples only a fraction of them. We use the following modeling for the sampling: each measurement of $\hat{N}_i(T_k)$ is a realization of a binomial distribution $B(d^k W_i^{T_k} N_i, p_k)$, where $d^k W_i^{T_k} N_i$ is the theoretical population of plasmids in the DNA sequencer carrying the mutation $i$ at time $T_k$, and $p_k$ is the inferred sampling rate of the

DNA sequencer at time $T_k$. We can estimate the sampling rate $p_k$ at different times

$$p_k = \frac{\sum_i \hat{N}_i(T_k)}{N_{OD}(T_k)} \ , \tag{3}$$

where $\sum_i \hat{N}_i(T_k)$ is the total population sampled with the DNA sequencer at time $T_k$, and $N_{OD}(T_k)$ is the theoretical population in the DNA sequencer at time $T_k$. The ratio of these two quantities is an estimation of the sampling rate of the DNA sequencer.

#### 1.2 Time parameterization

According to the measurement protocol, the different times steps $T_k$ correspond to a number of population-averaged generations ($T_0 = 0$, $T_1 = 4$, $T_2 = 9$, $T_4 = 19$ and $T_6 = 29$)[1]. We have the following evolution for total number of plasmids

$$N(T_k) = 2^{T_k - T_{k-1}} N(T_{k-1}) \ . \tag{4}$$

However, as a significant fraction of mutants may die due to the antibiotic, the number of generations may be underestimated during a cycle. Therefore, we redefined the time scale by the number of generations the wild-type did. Within this new definition of time $T^{WT}$, the absolute fitness of the wild-type $W_{WT}$ must be equal to 2. We found indeed this value with our inference procedure (Fig. 1).

Between $T_{k-1}$ and $T_k$, we expect to have a $T_k - T_{k-1}$ population-averaged doubling. During this cycle, the wild-type has made $T_k^{WT} - T_{k-1}^{WT}$ doubling. $\hat{N}^{WT}(T_k)$ is the measured population of wild-type at time $T_k$. If $f(k)$ is the frequency of wild-type after the $k^{\text{th}}$ cycle

$$f(k) = \frac{\hat{N}^{WT}(T_k)}{\sum_i \hat{N}_i(T_k)} \ , \tag{5}$$

as

$$2^{T_k^{WT} - T_{k-1}^{WT}} f(T_{k-1}) \sum_i \hat{N}_i(T_{k-1}) = f(T_k) \sum_i \hat{N}_i(T_k) \ , \tag{6}$$

we get,

[1]The dilution factor $d = \frac{1}{32} = \frac{1}{2^5}$ corresponds to 5 population-averaged doubling, which explains why $T_{k+1} - T_k = 5$, except for $T_1 - T_0 = 4$, as at $T_0$ $OD_{600} = 0.4$, and therefore after the dilution only 4 population-averaged doubling are needed to reach $OD_{600} = 0.2$.

$$T_k^{WT} - T_{k-1}^{WT} = T_k - T_{k-1} + \log_2 \frac{f(T_k)}{f(T_{k-1})} \ . \tag{7}$$

And then

$$T_k^{WT} = \sum_{k'=1}^{k} (T_{k'} - T_{k'-1}) + \log_2 \frac{f(T_{k'})}{f(T_{k'-1})} \tag{8}$$

$$= T_k + \log_2 \frac{f(k)}{f(0)} \ . \tag{9}$$

The results are reported in Table 1.

| | k = 0 | k = 1 | k = 2 | k = 4 | k = 6 |
|---|---|---|---|---|---|
| $T_k$ | 0 | 4 | 9 | 19 | 29 |
| $T_k^{WT}$ | 0 | 6.6 | 11.9 | 22.0 | 32.1 |

Table 1 – Comparison between $T_k$ and $T_k^{WT}$

In the following, we use $T_k^{WT}$ as reference time scale, but we note it as $T_k$ to simplify the notations.

#### 1.3 Computation of the likelihood

For a given mutation $i$, we want to estimate the absolute fitness $W_i$ knowing the measurements of the population $\{\hat{N}_i(T_k)\}_k$ at different times $T_k$. Within our model, the probability of $\{\hat{N}_i(T_k)\}_k$ at different times $T_k$ knowing $W_i$ can be written as

$$P(\{\hat{N}_i(T_k)\}_k | W_i) \propto \sum_{N_i} \prod_k \binom{d^k W_i^{T_k} N_i}{\hat{N}_i(T_k)} (1 - p_k)^{d^k W_i^{T_k} N_i - \hat{N}_i(T_k)} p_k^{\hat{N}_i(T_k)} \ , \tag{10}$$

Without a priori knowledge of the distribution of $W_i$, using Bayes' theorem, the likelihood can be written as

$$P(W_i | \{\hat{N}_i(T_k)\}_k) \propto P(\{\hat{N}_i(T_k)\}_k | W_i) \ . \tag{11}$$

The likelihood for all the mutations reads

$$P_{\text{model}}(W_i | \{\hat{N}_i(T_k)\}_k) \propto \prod_i P(W_i | \{\hat{N}_i(T_k)\}_k) \ . \tag{12}$$

In the most general case, because of the binomial coefficients in equation 10, the exact likelihood can not be computed analytically or numerically. Nonetheless, we can use a Gaussian approximation for the likelihood. Within this approximation, we have

$$P_{\mathrm{model}}(W_i|\{\hat{N}_i(T_k)\}_k) \quad \propto \quad \max_{\{N_i\}} \exp\left(\Phi(\{W_i, \hat{N}_i(T_k), N_i\}) - \frac{1}{2}\log|\Phi^{''}(\{W_i, \hat{N}_i(T_k), N_i\})|\right) \ ,$$

where

$$\begin{aligned}
\Phi(\{W_i, \hat{N}_i(T_k), N_i\}) \quad &= \quad \sum_i \sum_k \log\,(d^k W_i^{T_k} N_i)(\frac{1}{2} + W_i^{T_k} N_i) \qquad (13) \\
&- \quad \log\,(d^k W_i^{T_k} N_i - \hat{N}_i(T_k))(\frac{1}{2} + d^k W_i^{T_k} N_i - \hat{N}_i(T_k)) \\
&- \quad \log\,(\hat{N}_i(T_k))(\frac{1}{2} + \hat{N}_i(T_k)) \\
&+ \quad \log\,(1 - p_k)(d^k W_i^{T_k} N_i - \hat{N}_i(T_k)) + \log\,(p_k)\hat{N}_i(T_k) \ .
\end{aligned}$$

The likelihood is maximized numerically with respect to $W_i$ and $N_i$. In some cases, we can compute exactly the true likelihood, and our Gaussian approximation is in good agreement with the true likelihood (Fig. 1).

Once the parameters have been inferred, we have the following log-likelihood

$$LL(\{W_i, \hat{N}_i(T_k), N_i\}) = \Phi(\{W_i, \hat{N}_i(T_k), N_i\}) - \frac{1}{2}\log|\Phi^{''}(\{W_i, \hat{N}_i(T_k), N_i\})| \ . \qquad (14)$$

Therefore, we can estimate the uncertainty on the parameter $W_i$ as

$$\sigma_{W_i} = \sqrt{\frac{1}{\frac{\partial^2 LL(\{W_i, \hat{N}_i(T_k), N_i\})}{\partial W_i^2}}} \ . \qquad (15)$$

As by definition of the time, $W_{WT} = 2$, and $\frac{\sigma_{W_i}}{W_i} \ll 1$, the standard deviation $\sigma_{\log(w_i)}$ associated with the relative fitness is equal to $\frac{\sigma_{W_i}}{W_i}$. In practice, we estimate log-fitness only between $T_0$ and $T_2$, because at longer times, as yet not understood effects seem to disrupt the exponential growth of bacteria.

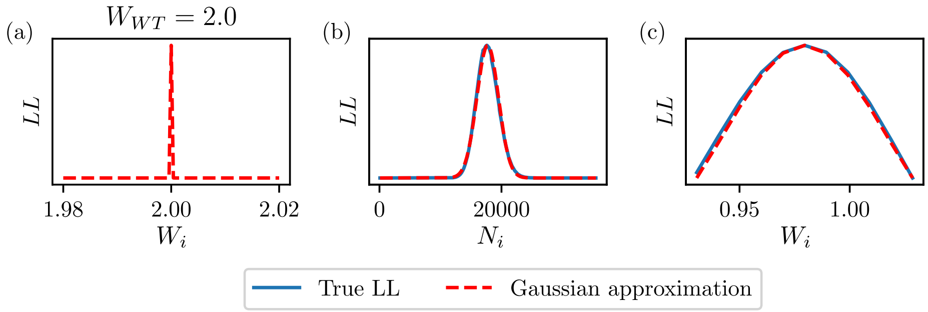


Figure 1 – (a). Estimation of $W_{WT}$. $W_{WT} = 2$ in agreement with the definition of the $T_k$. (b) and (c). Comparison between the exact log-likelihood and the Gaussian approximation for a given mutant. (b) Comparison between the exact log-likelihood and the Gaussian approximation for the estimation of $N_i$. (c) Comparison between the exact log-likelihood and the Gaussian approximation for the estimation of $W_i$.

#### 1.4 Consistency with Minimum Inhibitory Concentrations and replicas

We can compare our estimations of the log-fitness with measures of Minimum Inhibitory Concentrations (MIC) through experiments at 1, 2, 4, 8, and 16 g/L of amoxicillin. MIC and log-fitness are very high correlated for both single (Spearman correlation, $\rho = 0.98$) and double mutants ($\rho = 0.76$), although in a non-linear way (Figure supplementary).

Furthermore, we performed a second experiment to measure again the log-fitness (a second replicate) and for both replicates, the log-fitness of the single mutants, double mutants, and epistasis are highly correlated (respectively, $r^2$=1.0, $r^2$=0.95, $r^2$=0.99, Figure supplementary).

In addition, the uncertainty of the log-fitness inferred by our inference procedure as well as that estimated from the two replicates is correlated ($\rho = 0.64$ for the single mutants, $\rho = 0.58$ for the double mutants, Fig. 2).

#### 1.5 Definition of the lethality threshold

Mutants were considered lethal under a theoretical lower threshold for log-fitness equals to $-\log(2)$. At this specific value, bacteria do not grow in the solution. In practice, to limit the noise, we used a higher threshold equals to $-0.6$.

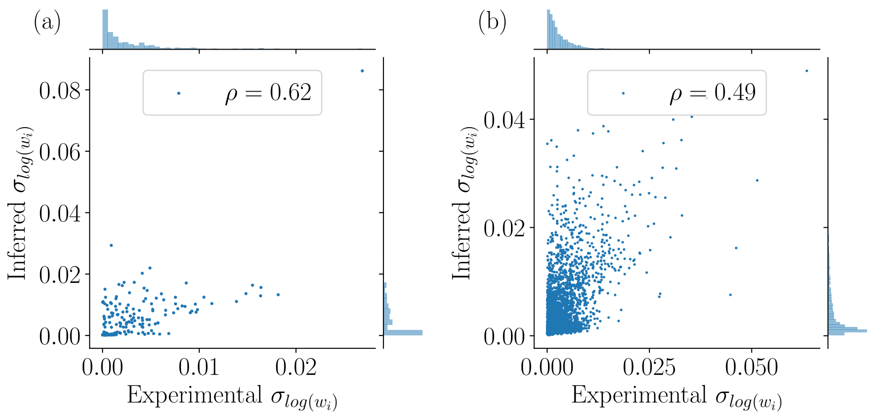


Figure 2 – Comparison between the experimental error and the inferred error of the log-fitness. (a) For single mutants. (b) For double mutants.

#### 2 Inference of the stability model

We denote as $w_i(a)$ the relative fitness of the mutant that as the amino acid $a$ at site $i$ of the $\alpha$-helix. We denote as $w_{i,j}(a,b)$ the relative fitness of the mutant that as the amino acid $a$ at site $i$ of the alpha-helix and amino acid $b$ at site $j$.

The stability model reads:

$$\log(\hat{w}_i^a) = \log\left(1+\exp\left(\frac{\Delta G_0}{RT}\right)\right) - \log\left(1+\exp\left(\frac{\Delta G_0 + \Delta\Delta G_i^a}{RT}\right)\right), \quad (16)$$

$$\log(\hat{w}_{i,j}^{a,b}) = \log\left(1+\exp\left(\frac{\Delta G_0}{RT}\right)\right) - \log\left(1+\exp\left(\frac{\Delta G_0 + \Delta\Delta G_i^a + \Delta\Delta G_j^b}{RT}\right)\right) \quad (17)$$

To fit the parameters of the thermodynamics model of stability parameters, we had to assign every single mutant a free entropy value, $\Delta\Delta G$, reflecting the impact of the mutant on the whole protein stability and an overall stability of the protein $\Delta G_0$. Though measures of $\Delta G_0$ have been done *in vitro*, the cellular environment in which the mutants are evaluated could substantially affect the value. We, therefore, have also estimated $\Delta G_0$. Ideally, estimated log-fitness is directly connected to $\Delta\Delta G$ for the single mutants, by inverting the equation 16, but is limited, as explained in the main text.

For the inference of the $\Delta\Delta G$ we keep only the single mutant with a log-fitness greater than $-0.6$. For each pair of previously chosen single mutants, the associated double mutant is kept if it exists. Its relative log-fitness is thresholded at $-0.6$. The stability model is itself thresholded at $-0.6$ during the inference.

The model is over-constrained. By defining the residue associated with a given mutation

$$r_i = \frac{\log(w_i) - \log(\hat{w}_i)}{\sigma_{\log(w_i)}} \ , \tag{18}$$

with the following cost function

$$C(\Delta G_0, \{\Delta\Delta G_i^a\}) = \frac{1}{2}\sum_i \alpha_i T^2 \Phi(\frac{r_i^2}{T^2}) \ . \tag{19}$$

$\alpha_i$ is a statistical weight. For the double mutants, $\alpha_i = 2$. For the single mutants, $\alpha_i$ is equal to the number of double mutants with this single mutation. The weighting gives an equivalent weight for the single mutants and the double mutants.

Here, we use $\Phi(x) = \arctan(x)$ in order to penalize the strong outliers. $T$ is a threshold that controls the importance of the regularization of the outliers and is chosen such that 30% of the mutations are considered as outliers. The results are consistent for a wide range of thresholds $T$.

### 3 Estimation of the error part of the stability model prediction

Using the whole data set, we could estimate an error to the model using a maximum likelihood framework. $\Delta\Delta G$ values were fixed. We estimated that the deviation of the observed log-fitness to the one predicted with the stability model resulted from an overall random deviation from the model. This deviation to the model could be either the same for all pairs of mutations or could be different for residues in contact or not, *i.e.* a two model parameters. The two errors model was always much better than the single error, and always suggested a higher deviation to the model for residues in contact compared to distant residues.

For the single error model:

$$\sigma_m = \sqrt{N^{-1} \sum_{i,j,a,b} \left(\log(w_{i,j}(a,b)) - \log(\hat{w}_{i,j}^{a,b})\right)^2} \ , \tag{20}$$

where $N^{-1}$ is the number of terms in the previous sum.

For the double error model:

$$\sigma_{md} = \sqrt{N_d^{-1} \sum_{i,j,a,b} (1-\delta_{i,j}) \left(\log(w_{i,j}(a,b)) - \log(\hat{w}_{i,j}^{a,b})\right)^2} \ , \tag{21}$$

$$\sigma_{mn} = \sqrt{N_n^{-1} \sum_{i,j,a,b} \delta_{i,j} \left(\log(w_{i,j}(a,b)) - \log(\hat{w}_{i,j}^{a,b})\right)^2} \ , \tag{22}$$

with $\delta_{i,j}$ if the chains of residues carrying mutation i and j are less than 6Å away and 0 otherwise. $N_d = \sum_{i,j,a,b} (1-\delta_{i,j})$ and $N_n = \sum_{i,j,a,b} \delta_{i,j}$, $(N = N_d + N_n)$.

We used also a model with a specific error between two given sites:

$$\sigma_{i,j} = \sqrt{N_{i,j}^{-1} \sum_{a,b} \left(\log(w_{i,j}(a,b)) - \log(\hat{w}_{i,j}^{a,b})\right)^2}\ , \tag{23}$$

where $N_{i,j}$ is the number of double mutations between the sites $i$ and $j$.

### 4 Computation of the p-value

If we choose as a null hypothesis that there is no relationship between pairs of sites with the largest idiosyncratic epistasis and pairs of sites with the largest Frobenius norm, the probability that among $L$ pairs of sites with the largest idiosyncratic epistasis, $c$ are also present in the $L$ pairs of sites with the largest Frobenius norm follows an hypergeometric distribution with parameters $N$, $L$ and $c$, where $N$ is the total number of possible pairs. Therefore, the p-value reads

$$p = \sum_{k=c}^{L} \frac{\binom{L}{k}\binom{N-k}{L-k}}{\binom{N}{L}} \tag{24}$$
